# DEDuCT 3.0: An enhanced and expanded FAIR-compliant resource and toxicology knowledge graph for endocrine disrupting chemicals

**DOI:** 10.64898/2026.01.23.701267

**Authors:** Nikhil Chivukula, Shrish Vashishth, Pavithra Kandasamy, Shreyes Rajan Madgaonkar, Areejit Samal

## Abstract

Endocrine disrupting chemicals (EDCs) are of particular regulatory and research interest due to the increasing incidence of endocrine-related disorders, such as declining fertility rates and reproductive health problems. The <u>D</u>atabase of <u>E</u>ndocrine <u>D</u>isr<u>u</u>pting <u>C</u>hemicals and their <u>T</u>oxicity Profiles (DEDuCT) has gained importance in both academic and regulatory settings by systematically curating data from published literature to characterize these chemicals. Given the growing body of EDC literature, this study aimed to consolidate the latest research and update this critical database. First, more than 14000 research articles were screened through an extensive four-stage manual process, and integrated with the earlier version to create the updated DEDuCTv3.0, comprising 1043 unique EDCs and 796 unique endocrine-related endpoints curated from 3269 published articles. Thereafter, human- and rodent-specific biological endpoint data including interacting genes/proteins, phenotypes, diseases, and adverse outcome pathways (AOPs) were curated from toxicology-relevant databases and systematically integrated with DEDuCTv3.0 to construct a large-scale toxicology knowledge graph for EDCs, termed DEDuCT-KG. DEDuCT-KG was then hosted on a Neo4j database and made easily accessible through a novel interactive user interface. The utility of DEDuCT-KG was demonstrated by exploring potential mechanisms of action associated with obesogenic EDCs within DEDuCTv3.0. Furthermore, the constructed EDC-AOP network, linking 949 EDCs to 381 AOPs within AOP-Wiki, revealed diverse toxicity mechanisms associated with EDCs. Integration with consumer product database and regulatory chemical lists showed that some of these EDCs are present in food contact materials, personal care products, and daily use items, highlighting potential exposure pathways. Overall, all data compiled in this study have been integrated into the DEDuCT webserver, which has been further enhanced to align with FAIR principles. In sum, this study provides a much-needed update to DEDuCT and offers a single point of access to EDC-relevant data to accelerate research and regulation of EDCs.

## 1. Introduction

Endocrine disrupting chemicals (EDCs) are of particular concern to human and environmental health, as they can interfere with the normal functioning of the endocrine system and lead to a variety of adverse effects, including reproductive, physiological, and developmental outcomes, among others.^1–3^ Growing evidence of declining fertility rates and reproductive health problems has increased concern about EDCs and has led to more studies and regulations across global bodies to mitigate their effects.^2,3^ In response to this need, the <u>D</u>atabase of <u>E</u>ndocrine <u>D</u>isr<u>u</u>pting <u>C</u>hemicals and their <u>T</u>oxicity profiles (DEDuCT) has emerged as a resource that organizes published evidence and makes scientific data on EDCs easily accessible for better understanding.^4,5^ The French Agency for Food, Environmental and Occupational Health & Safety (ANSES) has recognized the DEDuCT’s four-stage criteria for extracting information from published literature as an important tool to gain knowledge on EDCs from published studies, and moreover, has used the data for designing their Second French National Strategy for Endocrine disruptors (SNPE2).^6^ Furthermore, DEDuCT has also been instrumental in providing a curated dataset of chemicals for the development of *in silico* methods for the prediction of EDCs.^7,8^ Moreover, the curated endocrine-mediated endpoint data within DEDuCT has been extensively integrated with the Adverse Outcome Pathway (AOP) framework to explore chemical-induced toxicity mechanisms across different classes of chemicals.^9,10^ In short, DEDuCT is an important resource not only for academic purposes but also one that has gained regulatory relevance.

Since the release of the previous version of DEDuCT in 2020, a large body of literature on EDCs has been generated. In parallel, while toxicologically-relevant databases, such as the United States Environmental Protection Agency’s (US EPA) ToxCast^11,12^ and CompTox Chemicals Dashboard,^13^ the Comparative Toxicogenomics Database (CTD),^14^ and AOP-Wiki (https://aopwiki.org), have been continuously updated and expanded, they do not specifically focus on the unique mechanisms and endpoints associated with endocrine disruptors. Although the previous version of DEDuCT included ToxCast data, it lacked integration with information from other resources and did not capture chemical structure information from the CompTox Dashboard. Therefore, to provide a single point of access to all EDC-related data and align the DEDuCT knowledgebase with Findable, Accessible, Interoperable and Reusable (FAIR) principles,^15^ there is a need to curate data from recently published literature and integrate it with data from these external resources.

In recent years, knowledge graphs have emerged as a framework to organize diverse information and effectively represent the complex interconnections across disparate data sources.^16–18^ In knowledge graphs, nodes represent entities of interest, and the edges denote the semantic relations that characterize the connections between the entities.^16–18^ Such structured representations have provided a basis for efficiently processing heterogeneous knowledge and have eventually supported the training and reasoning of modern artificial intelligence (AI) systems.^16–18^ In particular, toxicology knowledge graphs have emerged as a tool to systematically integrate existing chemical-associated toxicity and biological information, enabling the discovery and expansion of knowledge on chemical-induced toxicities.^19^ For instance, Evangelista *et al*.^19^ organized chemical-specific genetic changes and reproductive disruptions into a knowledge graph representation, enabling the exploration of the mechanisms of action of previously overlooked reproductive toxicants. While there are other toxicological knowledge graphs, such as TOXIN-KG^20^ which particularly focus on cosmetic-induced liver toxicities, and FORUM^21^ and AidTox^22^ that focus on drug-induced toxicities, there is no large-scale knowledge graph that captures EDC-specific toxicological information. In this direction, the compiled and curated data in DEDuCT can be organized to construct such a large-scale toxicological knowledge graph for EDCs and aid in providing insights into EDC-associated toxicities.

In this study, more than 14000 articles were first identified as the new published literature that had not been screened in the previous versions of DEDuCT. These articles were then manually screened using the established four-stage workflow. Thereafter, the associated adverse effects were assigned to standardized endocrine-mediated endpoints and mapped to systems-level perturbation events. All these compiled data were then integrated with existing data to create the latest version 3.0, referred to as DEDuCTv3.0 in this study. The associated human- and rodent-specific gene targets were then curated from ToxCast invitrodb v4.3 and CTD, along with associated phenotype and disease information from CTD. The AOPs were curated from AOP-Wiki, and the EDCs were systematically mapped to these AOPs to explore diverse toxicity mechanisms associated with EDCs. All the compiled data were then integrated to construct a large-scale toxicology knowledge graph for EDCs. This knowledge graph has been made accessible to the research community through an interactive user interface on the DEDuCT webserver. The utility of the constructed knowledge graph was then demonstrated by identifying potential mechanisms of action associated with obesogenic EDCs within DEDuCTv3.0. Furthermore, consumer product data was accessed from the US EPA’s Chemical and Products Database and used to explore potential exposure pathways. Several chemical lists were used to explore regulatory coverage. All the curated data, including the novel toxicological knowledge graph, has been made accessible on the enhanced and FAIR-compliant DEDuCTv3.0 webserver available at: https://cb.imsc.res.in/deduct/. In sum, this study presents a much-needed update to the resource and, most importantly, organizes diverse toxicological data as a knowledge graph to enable future research and aid in better chemical regulation strategies for EDCs.

## 2. Results and Discussion

### 2.1. DEDuCTv3.0 - an enhanced resource on potential endocrine disrupting chemicals

The <u>D</u>atabase of <u>E</u>ndocrine <u>D</u>isr<u>u</u>pting <u>C</u>hemicals and their <u>T</u>oxicity profiles (DEDuCT) is a unique repository of endocrine disrupting chemicals (EDCs), and their associated endocrine-mediated effects that were systematically curated from published literature.^4,5^ Since the release of DEDuCT version 2.0 in October 2020, there has been substantial increase in literature focusing on characterization of EDCs, with PubMed now cataloging more than 14000 EDC-associated articles that were not previously screened (Supporting Information; Figure 1). This is the primary motivation to update and release the latest version, DEDuCT version 3.0 (DEDuCTv3.0).

**Figure 1.**
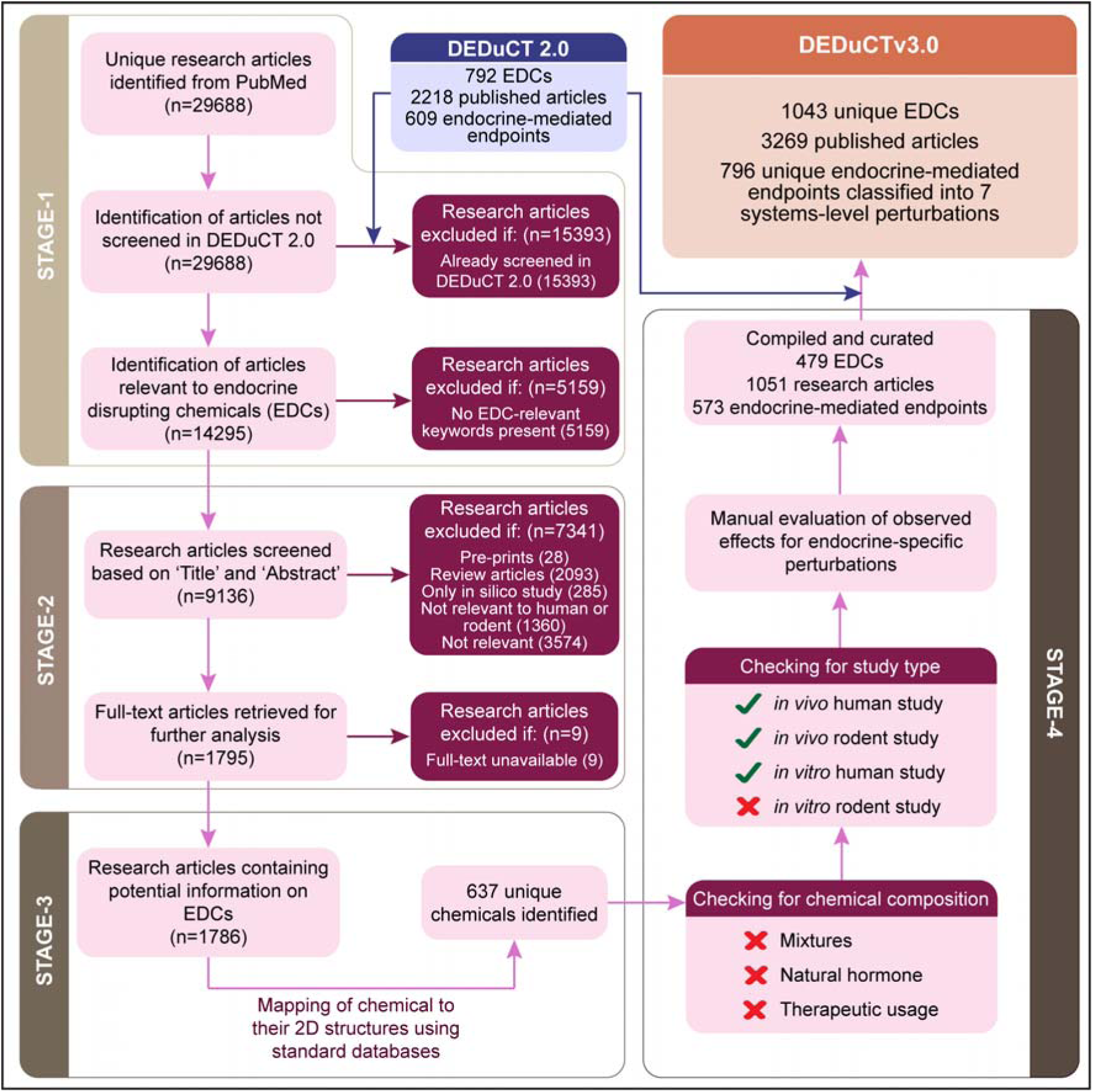
The workflow to identify published literature containing endocrine disrupting chemicals and their associated endpoints is presented as a PRISMA statement.

Here, we present DEDuCTv3.0, cataloging 1043 unique potential EDCs, systematically curated from 3269 published articles (Methods; Figure 1; Table S1). These chemicals were linked to 796 unique endocrine-mediated endpoints, which were further classified into seven functional categories (Supporting Information; Table S2). It was observed that most endpoints were categorized as ‘Reproductive endocrine-mediated perturbations’ (Supporting information; Figure S1a). Moreover, it was also observed that 823 of 1043 EDCs were associated with endpoints categorized as ‘Reproductive endocrine-mediated perturbations’ (Figure S1b). Additionally, based on the previous study,^4^ the standardized dosage information was utilized to compute the No Observed Adverse Effect Level (NOAEL) and Low Observed Adverse Effect Level (LOAEL) to enable further quantitative assessments of EDC-associated toxicities (Table S3).

Notably, in comparison to previous version, DEDuCTv3.0 includes 251 new chemicals, and 187 new endocrine-mediated endpoints, demonstrating a significant expansion of coverage of chemicals and their associated biological effects. Furthermore, 150 of these 251 new chemicals have reported evidence from *in vitro* human tests, suggesting a methodological shift toward more efficient, human-relevant toxicity testing of potential EDCs (Table S1). Such shifts are consistent with the framework of New Approach Methodologies (NAMs) and underscore their utility in streamlining the identification of potential EDCs.^3^ In addition, compared with other EDC-relevant data compilations, such as World Health Organization (WHO) report,^23^ the Endocrine Disruption Exchange (TEDx; https://endocrinedisruption.org/) and EDCs Databank version 2015,^24^ DEDuCTv3.0 now compiles 435 EDCs not captured by any of these resources (Figure S2a). Importantly, among these 435 EDCs, 193 were newly compiled in DEDuCTv3.0 (Figure S2b), emphasizing the necessity for this effort to enable further research in EDCs.

An environmental source-based classification of EDCs in DEDuCTv3.0 revealed that 466 chemicals originated from ‘Industry’, while 461 chemicals originated from ‘Consumer products’ (Supporting Information; Table S1). Next, based on the hierarchical structural classification of EDCs provided by ClassyFire,^25^ it was observed that 988 EDCs were classified as organic compounds, of which 242 chemicals were newly compiled in DEDuCTv3.0, while 55 were classified as inorganic compounds, of which 9 were newly compiled in DEDuCTv3.0 (Supporting Information; Table S1).

Furthermore, based on the previous study,^4^ the chemical space within DEDuCTv3.0 was first visualized as a chemical similarity network (CSN), that was constructed with 1043 EDCs as nodes and the structural similarity > 0.5 as edges (Supporting Information; Figure S3). It was observed that there were 99 connected components containing at least 2 nodes, of which the largest connected component comprised 248 nodes, while there were 409 isolated nodes (Figure S3). Notably, 90 of the 251 new chemicals in DEDuCTv3.0 were present as isolated nodes, suggesting they are structurally distinct from the chemicals previously compiled (Figure S3). Moreover, 145 of the 251 chemicals were structurally similar to chemicals compiled in DEDuCT version 2.0, while 16 others were structurally distinct (Figure S3). Subsequently, RDKit was utilized to compute chemical scaffolds according to Bemis-Murcko definition.^26^ It was observed that 813 of 1043 EDCs share 323 unique scaffolds, among which 75 scaffolds are shared by at least 2 chemicals (Table S1; Figure 2). Additionally, the chemical space was visualized as a target similarity network (TSN), that was constructed with 579 of 1043 EDCs with human- or rodent-relevant target gene information within ToxCast invitrodb v4.3.^12^ In this TSN, the 579 chemicals were considered as nodes, and the target similarity > 0.5 were considered as edges (Supporting Information; Figure S4). It was observed that there were 34 connected components containing at least 2 nodes, of which the largest connected component comprised 76 nodes, while there were 408 isolated nodes (Figure S4). This dissimilarity in EDC-associated gene targets suggests that these EDCs trigger diverse molecular initiating events. Moreover, based on the previous study,^4^ a lack of correlation was observed between the chemical and target similarity of the 579 of 1043 potential EDCs within DEDuCTv3.0 (Supporting Information; Figure S5).

**Figure 2.**
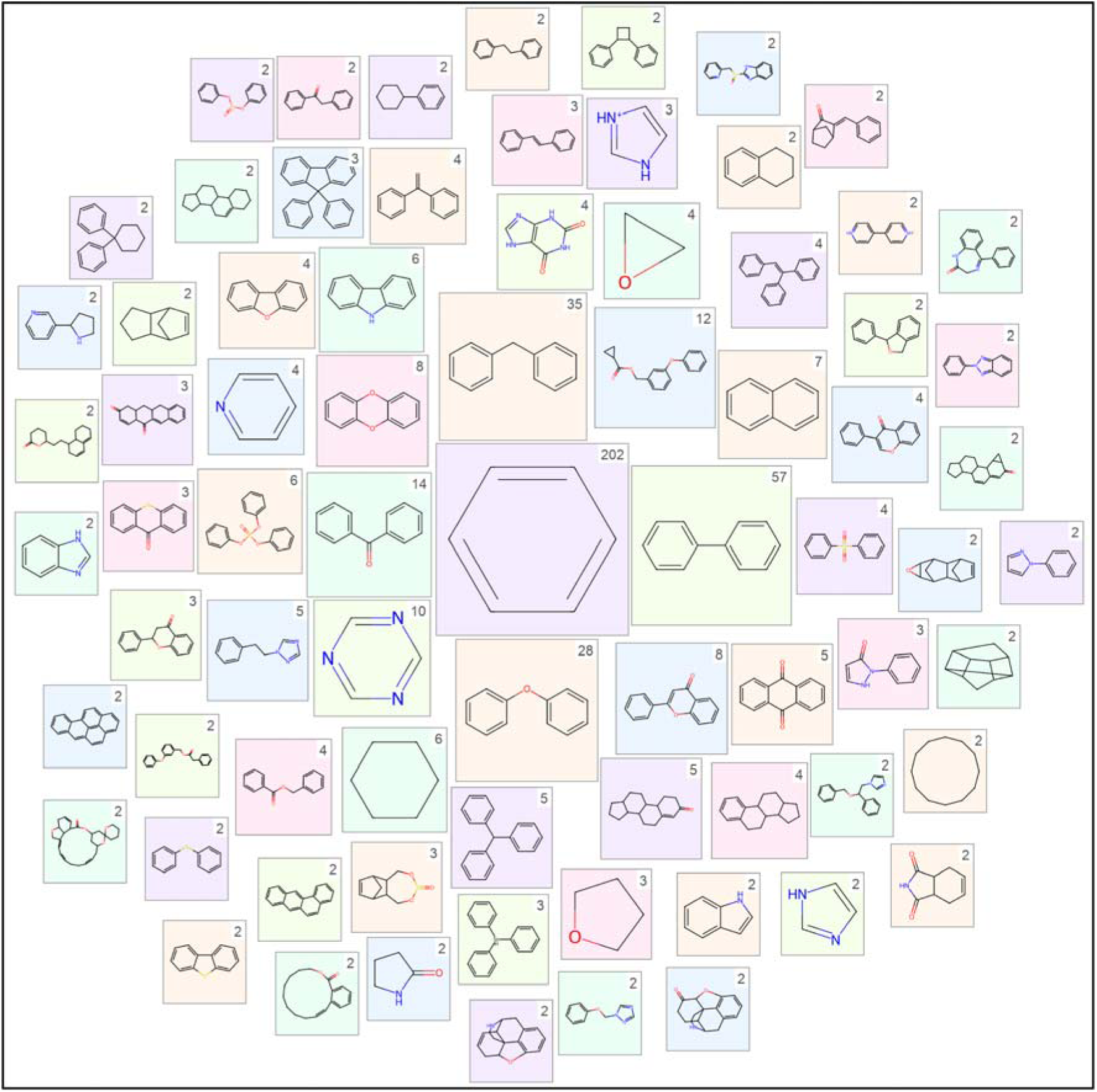
A scaffold cloud representation of the 75 unique scaffolds that are shared between at least two chemicals. The scaffolds are computed using RDKit based on Bemis-Murcko definition. The number of chemicals sharing the scaffold is mentioned at the top-right corner of each box containing the scaffold.

Overall, DEDuCTv3.0 provides the most up-to-date and comprehensive information on potential EDCs and supports further research in systems toxicology and human exposome. Compared to previous versions, DEDuCTv3.0 further organizes the compiled data as a novel toxicology knowledge graph for EDCs, thereby enhancing the utility of the resource for knowledge generation in EDC research.

### 2.2. DEDuCT-KG: A toxicology knowledge graph on EDCs

A toxicology knowledge graph organizes chemical-relevant toxicological data from diverse sources into a network comprising chemical-associated toxicological data connected through semantic relationships.^19^ Such a construction enables the querying of subgraphs that can highlight previously unexplored associations between entities, thereby facilitating novel understanding of chemical-induced toxicities.^19^ In this study, the EDCs and their associated endocrine-mediated endpoints from DEDuCTv3.0 were utilized to construct a large-scale toxicology knowledge graph comprising 75106 nodes and 1065949 edges, and referred to as DEDuCT-KG (Supporting Information; Figure 3). Among these nodes, 1043 are EDCs, 796 are their associated endocrine-mediated endpoints from DEDuCTv3.0, 48999 are genes, 3098 are diseases, 19238 are phenotypes, 1547 are Key Events (KEs) within AOP-Wiki, and 385 are the curated AOPs from AOP-Wiki. Based on the association information from different sources, the edges were classified into 31 distinct edge types connecting these nodes. Moreover, the largest connected component (LCC) comprised 75048 nodes and 1065913 edges, which connected 1043 EDCs, 796 endpoints, 3096 diseases, 19219 phenotypes, 48979 genes, 1532 KEs, and 383 AOPs. This LCC signifies a vast corpus of information associated with the EDCs curated within DEDuCTv3.0. Moreover, DEDuCT-KG has been hosted on a Neo4j graph database (https://neo4j.com/) and made available on the webserver through an interactive user interface^27^ to enable further research (Methods; Figure 4).

**Figure 3.**
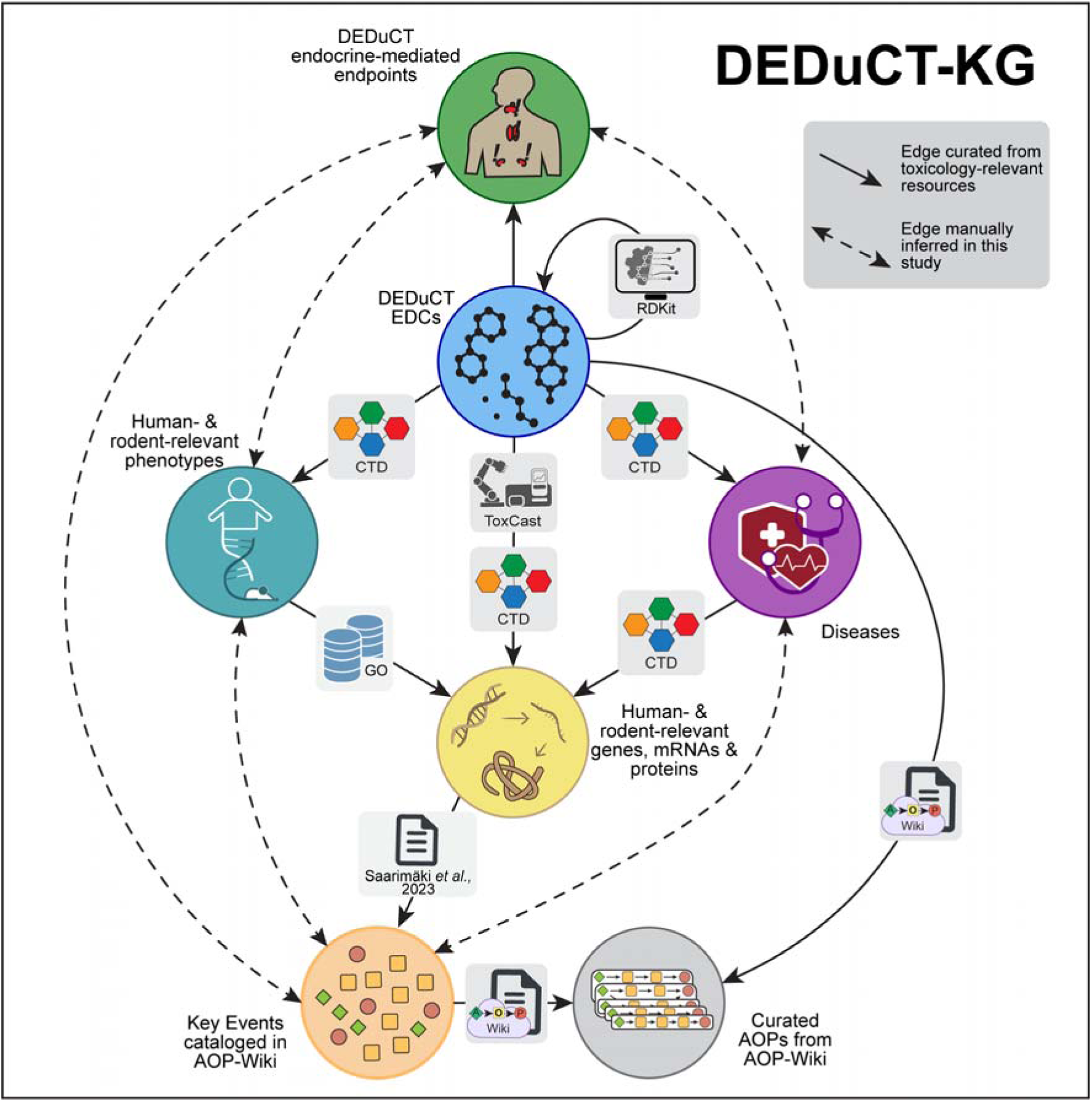
Network representation of different associations between 7 types of nodes present in DEDuCT Knowledge Graph (DEDuCT-KG). The resources from where each association was obtained is mentioned on the edge connecting the corresponding nodes.

**Figure 4.**
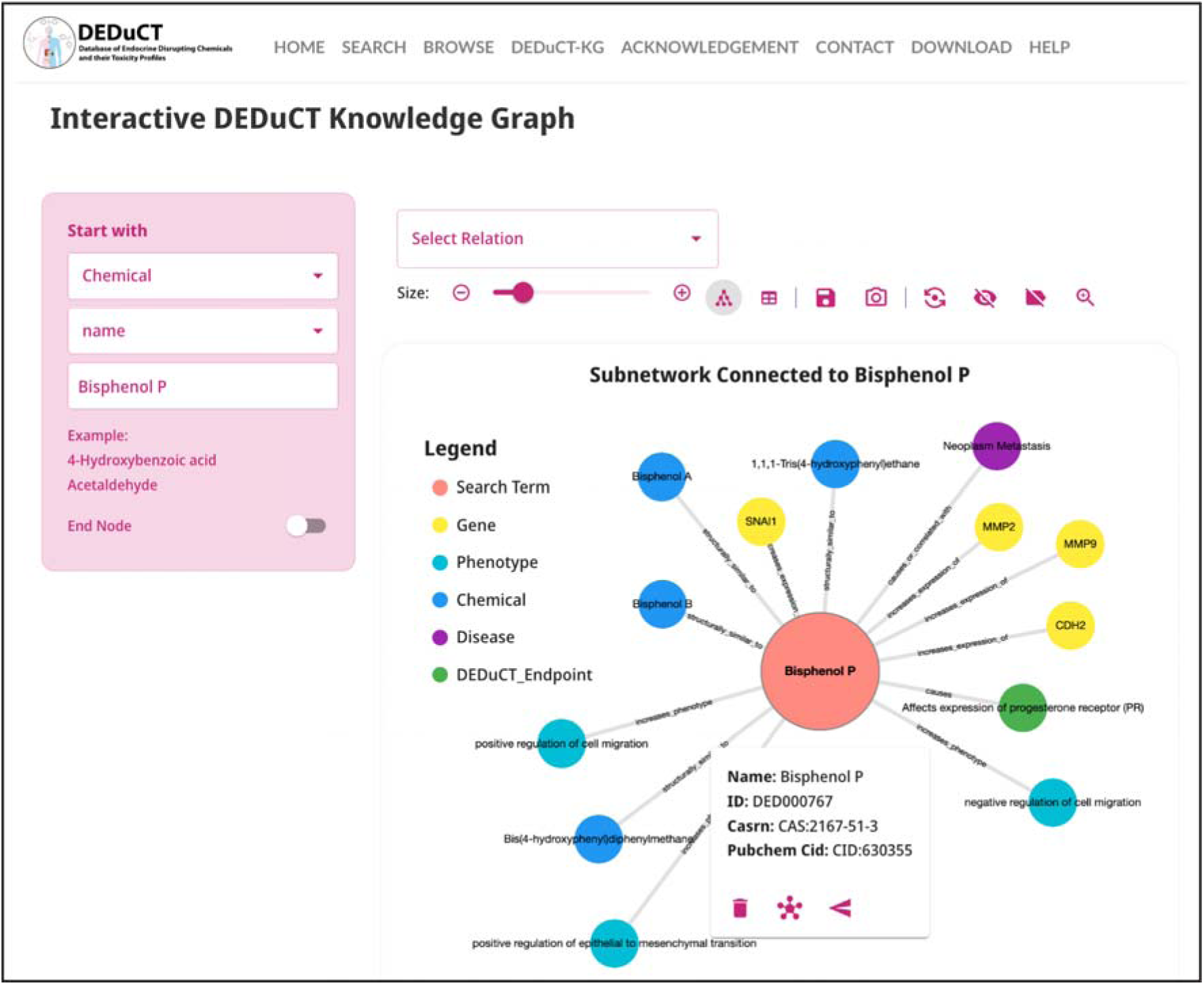
Interactive user interface in webserver of DEDuCTv3.0 for exploring the toxicology knowledge graph DEDuCT-KG.

In a previous study of reproductive toxicant chemicals, it was shown that knowledge graph construction enables the exploration of underexplored mechanisms of action (MOAs) associated with those chemicals.^19^ Similarly, in this study, the EDC and associated endocrine-mediated endpoint data were utilized to gain more information on EDC-associated MOAs. It was observed that the endocrine-mediated endpoints were not directly associated with the genes to help gauge their endocrine-specific MOAs (Figure 3). Instead, the endocrine-mediated endpoints were linked to phenotypes and diseases, which were further linked to genes (Figure 3). Previously, the concept of a Chemical-Gene-Phenotype-Disease (CGPD) tetramer was proposed as a bioinformatics-driven framework to bridge knowledge gaps between chemical exposures and outcomes by identifying intermediate molecular mechanisms.^28^ In this work, the framework was adapted by integrating endocrine-mediated endpoint-phenotype and endocrine-mediated endpoint-disease associations derived from the constructed knowledge graph (Supporting Information; Table S4). This allowed for the construction of subgraphs such as chemical-gene-phenotype-endpoint and chemical-gene-disease-endpoint, which revealed indirect gene-to-endpoint links. These genes can then be used to explore the potential MOAs associated with the chemical-induced endocrine-mediated effects.^19,28^ The utility of this knowledge graph is demonstrated using the following case study.

#### 2.2.1. Potential mechanism of action associated with obesogens within DEDuCTv3.0

Obesogens are a class of EDCs that promote obesity in exposed individuals by altering metabolic homeostasis and increasing susceptibility to weight gain.^29^ It was observed that DEDuCTv3.0 contained an endocrine-mediated endpoint titled ‘Lead to obesity’, which was categorized as ‘Metabolic endocrine-mediated perturbations’ (Table S2). Therefore, this endpoint was utilized to gain more insight into the potential obesogens within DEDuCTv3.0. From the constructed knowledge graph, this endpoint was observed to be associated with 52 chemicals, and diseases such as ‘Obesity, Abdominal’ (MESH:D056128), ‘Weight Gain’ (MESH:D015430), ‘Obesity’ (MESH:D009765), ‘Overweight’ (MESH:D009765), and ‘Obesity, Morbid’ (MESH:D009767), that were compiled from CTD (Supporting Information). Subsequently, the chemical-gene, chemical-disease and gene-disease links were selected from the knowledge graph to construct a chemical-gene-disease-endpoint subgraph comprising 18 chemicals, 260 genes, 4 diseases, and 1 endpoint (Table S4). Importantly, a prior study by Heindel *et al.*^29^ classified the chemical groups to which these 18 compounds belong as obesogenic, thereby supporting the construction of our knowledge graph and highlighting its utility in extracting toxicity relevant information.

Among the 18 chemicals, bisphenol A (CAS:80-05-7) was linked to 256 out of 260 genes, followed by tetrachlorodibenzodioxin (CAS: 1746-01-6) linked to 244 genes (Table S5). Among the 260 genes, TNF (NCBI Gene Id: 7124) and AHR (NCBI Gene Id: 196) were linked to all 18 chemicals, followed by NFE2L2 (NCBI Gene Id: 4780), CD36 (NCBI Gene Id: 948) and ADIPOQ (NCBI Gene Id: 9370) which were linked to at least 15 of the 18 chemicals (Table S5). Alternatively, PHAROS (https://pharos.nih.gov/) is an NIH-based repository that integrates data from the Target Central Resource Database (TCRD) to enable gene-disease association studies.^30^ From this database, 370 genes were found to be associated with ‘obesity disorder’, including the highly connected TNF, AHR, CD36, and ADIPOQ genes. The overlap of genes between the knowledge graph and PHAROS underscores the biological relevance of these targets. For instance, the overexpression of TNF gene and its role in inducing inflammation have been well-studied as central factors in the pathogenesis of obesity.^31–33^ AHR gene is found to have a dual role in pathogenesis of obesity,^34^ with its activation in adipocytes promoting their differentiation and eventually leading to promotion of obesity.^35^ The CD36 gene, that functions as a fatty acid translocase, showed increased expression in adipocytes of obese mice,^36^ and positively correlated with abdominal adipose tissue distribution in humans.^37^ Finally, the dysregulation of ADIPOQ gene has been extensively studied in pathogenesis of metabolic syndromes, including obesity.^38,39^

Next, chemical-gene-disease-endpoint subgraphs were selected for two obesogens, namely, dibutyl phthalate (DiBP; CAS:84-74-2) and perfluorooctane sulfonic acid (PFOS; CAS:1763-23-1), to further explore their MOAs (Figure 5). From these subgraphs, it was observed that these two chemicals are associated with TNF, AHR and CD36 genes (Figure 5). In particular, these chemicals were found to increase expression of TNF and CD36 genes based on evidence curated from CTD (Figure 5). Additionally, DiBP and PFOS were found to activate AHR, based on evidence from CTD and ToxCast invitrodb v4.3, respectively (Figure 5). These observations highlight that the two chemicals can potentially induce obesity through interaction with TNF, AHR, and CD36. Overall, this case study shows that the constructed DEDuCT-KG can be utilized to identify MOAs associated with EDCs within DEDuCTv3.0.

**Figure 5.**
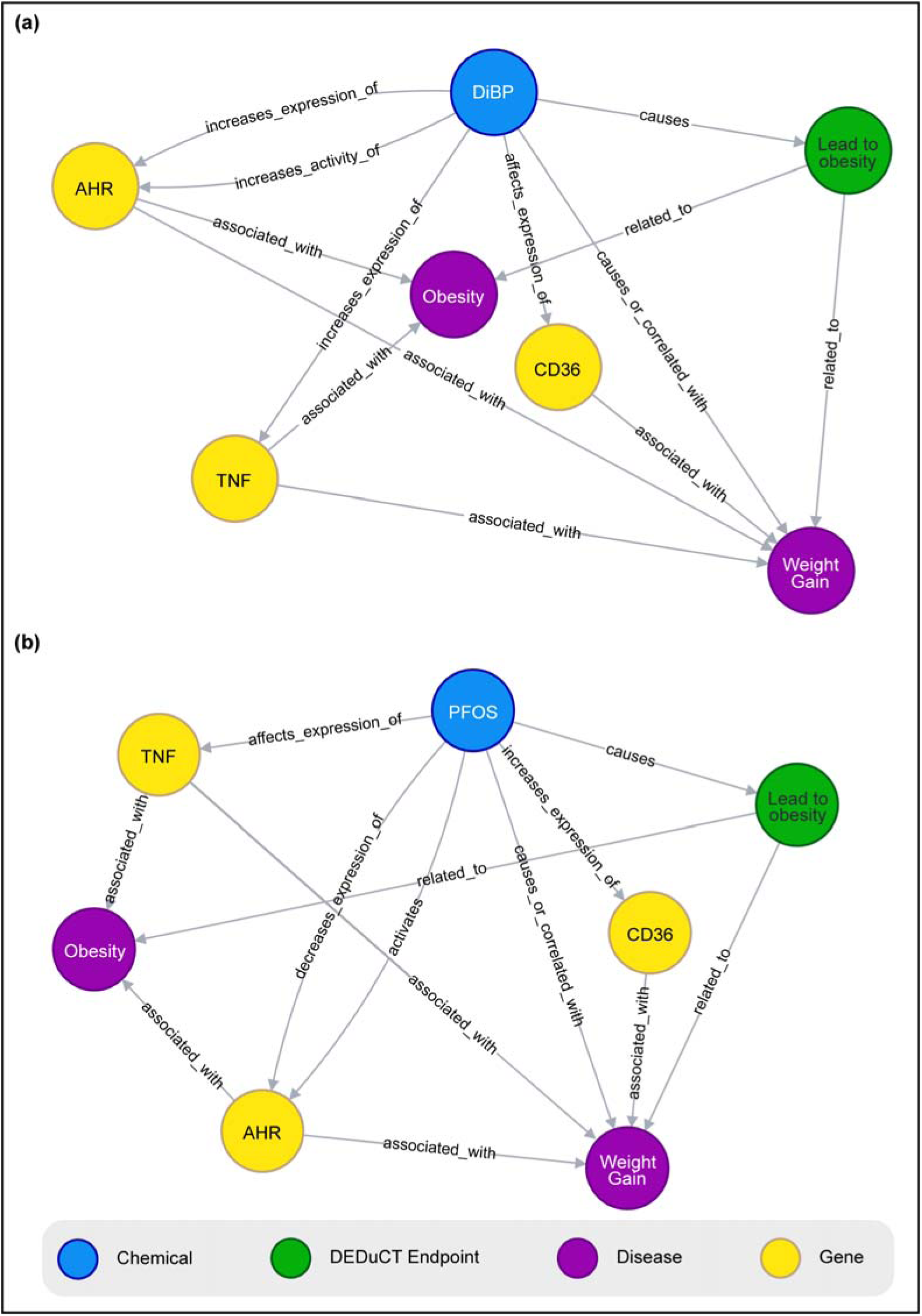
Chemical-gene-disease-endpoint subgraphs depicting the genes, diseases, and endocrine-mediated endpoint obtained from DEDuCT-KG, associated with obesogens: (**a**) dibutyl phthalate (DiBP; CAS:84-74-2); (**b**) perfluorooctane sulfonic acid (PFOS; CAS:1763-23-1). These subgraphs were generated using Neo4j.

### 2.3. Potential exposure to EDCs through consumer products

Humans are daily exposed to a wide variety of products, some of which may contain potential EDCs that pose a risk to human health. Therefore, the Chemical and Products database v4 (CPDat)^40^ was utilized to explore the exposure sources of potential EDCs in DEDuCTv3.0. First, 542 of 1043 EDCs were identified to be cataloged in CPDat, where the corresponding products were grouped into product use categories (PUC) (Table S1). It was observed that CPDat utilizes a hierarchical classification of PUCs, with the ‘Kind’ being the highest-level that describes the physical nature of the product.^40^ Specifically, ‘Article’ denotes products that are intended to be used for long-term use in the environment, ‘Formulation’ denotes products that are stored in containers, and ‘Occupation’ denotes products that are intended to be used in work environments.^40^

Among the 542 EDCs present in CPDat, 171 were identified as chemicals present in products categorized as ‘Articles’, where it was observed that 133 of 171 chemicals are present in products categorized as ‘Construction and building materials’, followed by 81 chemicals in products categorized as ‘Batteries’ (Table S1). This potentially suggests that such chemicals need to be checked for chronic or low-dose exposures. Similarly, 334 of 542 EDCs were identified as chemicals present in products categorized as ‘Formulation’ (Table S1). Among these chemicals, 168 are present in products categorized as ‘Personal care’, followed by 138 in ‘Home maintenance’ (Table S1). This highlights a significant pathway of exposure to potential EDCs through common consumer products. Finally, 418 of 542 EDCs were identified to be present in products categorized as ‘Occupational’ (Table S1). Among these 344 are present in products categorized as ‘Laboratory supplies’, followed by 198 chemicals as ‘Raw materials’ (Table S1).

In addition to PUC data, CPDat provides the functional uses associated with the chemicals in consumer products.^40^ This information is valuable for identifying less toxic chemical alternatives to replace the chemical of potential concern. To refine this dataset, chemicals with reported function as ‘unknown’ or ‘not reported’ were filtered out, resulting in the identification of 473 of 1043 EDCs with known functional use data (Table S1). Among these, it was observed that 417 chemicals were associated with 73 standardized Organisation for Economic Co-operation and Development (OECD) functional terms (Table S1).^40^ Analysis of these terms revealed that 179 chemicals function as ‘Biocides’, followed by 92 chemicals that function as ‘Fragrances’, in their corresponding products.

### 2.4. Distribution of EDCs in chemical lists

EDCs are of prime concern and are increasingly being regulated all around the world. To check the regulatory coverage of EDCs compiled in this study, several chemical lists were utilized. First, a unique list of 2049 EDCs was compiled from DEDuCTv3.0, the WHO report,^23^ the TEDx (https://endocrinedisruption.org/) and EDCs Databank version 2015.^24^ Next, 54 different chemical lists were compiled and categorized as substances in use and substances of concern (Methods; Table S6). The chemicals within each were categorized into three groups (groups I - III), with 19501 chemicals in group I, 11394 in group II, and 5488 in group III. An overlap with EDCs revealed 135 EDCs in group I, 581 in group II, and 353 in group III (Figure S6a). Among these, 20 are newly compiled within DEDuCTv3.0 in group I, 67 are newly compiled in group II, and 50 are newly compiled in group III (Figure S6a).

Next, the presence of these 3 groups of EDCs was checked in different categories of chemical lists. It was observed that the newly compiled chemicals were present in all the different categories, predominantly belonging to group II (Figure S6b). Notably, 15 of the new group I chemicals were identified to be currently in use as food additives or food contact materials, highlighting potential concern about dietary exposure. Among these 15 chemicals, 11 contained information from *in vitro* human cell line studies on evidence of endocrine-mediated effects (Table S1). Furthermore, 3 of the new chemicals were present in cosmetics, with 2 chemicals having *in vitro* human evidence of endocrine-mediated effects (Table S1). It was also observed that none of these new chemicals were cataloged as high production volume chemicals by either the US EPA or OECD. Overall, these findings highlight that some of the newly identified EDCs in DEDuCTv3.0 are currently in active use but remain unflagged by major regulatory agencies, demonstrating the utility of this compilation in uncovering previously overlooked chemicals of concern.

### 2.5. Exploration of EDC-relevant toxicity mechanisms through stressor-AOP network

EDC research is increasingly transitioning towards the adoption of NAMs.^3^ Within NAMs, the Adverse Outcome Pathways (AOPs) serve as an *in silico* toxicological framework for characterizing stressor-induced toxicity mechanisms and aiding in their risk assessment strategies.^3,41^ For DEDuCTv3.0, the stressor-AOP network was constructed by connecting 949 of 1043 EDCs and 381 curated AOPs from AOP-Wiki (https://aopwiki.org/), based on 759 associated Key Events (KEs) (Supporting Information; Tables S7-S10). Such a construction enabled the exploration of EDC-induced toxicity mechanisms through the AOP framework. It was observed that bisphenol A (CAS:80-05-7) had maximum associated KEs (390 KEs), which is expected given its extensive toxicological characterization within DEDuCTv3.0 (Table S10). It is immediately followed by cadmium (CAS:7440-43-9), which is linked to 289 KEs, and bis(2-ethylhexyl) phthalate (CAS:117-81-7), which is linked to 260 KEs (Table S10). Moreover, it was observed that the KE ‘Increased, Cancer’ (KE:885) was associated with highest number of EDCs, i.e., 348 of 943 EDCs, signifying that several chemicals have carcinogenic potential.

The EDC-AOP associations have been further classified based on levels of relevance, wherein Level 5 denotes that there is evidence of both molecular initiating event (MIE) and adverse outcome (AO) within an AOP associated with the chemical, and that the MIE and AO are connected through causal linkages (Supporting information).^10^ It was observed that 250 EDCs were associated with 215 curated AOPs from AOP-Wiki (Table S10). Notably, 96 EDCs were associated with 31 endorsed AOPs, signifying the utility of AOPs in characterizing EDC-specific mechanisms (Table S7). Moreover, 29 of the 215 AOPs are categorized as cancer based on their AOs, followed by 17 AOPs each for reproductive, gastrointestinal, and cardiovascular diseases (Table S7). Note that the current bias towards carcinogenic mechanisms may stem from the overrepresentation of cancer-associated AOPs within AOP-Wiki.^10^

### 2.6. Enhancements to DEDuCT webserver

The DEDuCT webserver, accessible at https://cb.imsc.res.in/deduct/, has been substantially updated to align more closely with the Findable, Accessible, Interoperable and Reproduceable (FAIR) principles^15^. Notably, the chemicals have been provided unique identifiers on DEDuCT, and are linked to their respective chemical pages in databases such as PubChem (https://pubchem.ncbi.nlm.nih.gov/) and Chemical Abstracts Service (CAS) Common Chemistry (https://commonchemistry.cas.org/), and the EPA’s CompTox Chemicals Dashboard^13^ (https://comptox.epa.gov/dashboard/) (Figure S7; Table S1). In this update, chemical pages have been enriched with more toxicology-relevant data curated from multiple relevant databases. For instance, information regarding chemical-associated genes, phenotypes, and diseases, has been curated from the CTD^14^ (Supporting Information) and made available as separate sections within the chemical pages (Figure S8). Additionally, the latest data from ToxCast invitrodb v4.3^12^ was systematically curated (Supporting Information) and made available in respective chemical pages for improved access (Figure S9). Here, the data is further categorized based on the type of target, such as RNA, protein, molecular messenger, pathway, cellular, and hormone. Moreover, the Product Use Categories (PUC) data and the functional uses of chemicals in consumer products have been curated from CPDat v4^40^ and made accessible through a dedicated section in the chemical page (Figure S10). The functional uses have been categorized under specific vocabularies from the US EPA and OECD (Figure S10), while the PUCs have been categorized based on the physical nature of the corresponding product (Figure S10). Regulatory lists were updated to include the latest chemical inventories used for chemical identification. Chemicals are searchable based on their identifiers, names, synonyms, physicochemical properties, or structures (Figure S11). Additionally, a large-scale toxicology knowledge graph (KG) for EDCs, DEDuCT-KG, was constructed and made accessible through an interactive Knowledge Graph UI^27^ on the webserver. Overall, DEDuCTv3.0 is a culmination of significant updates to the resource, making it more FAIR-compliant and enhancing the toxicological knowledge associated with the compiled EDCs. The data compiled in DEDuCTv3.0 has been made accessible through machine-readable file formats in the Downloads section of the webserver, enabling further research on EDCs.

## 3. Methods

### 3.1. Workflow to identify endocrine disrupting chemicals

Database of Endocrine Disrupting Chemicals and their Toxicity profiles (DEDuCT) is one of the largest resources on EDCs, that systematically compiles chemical-associated toxicities via extensive manual curation from published literature.^4,5^ Since the release of DEDuCT version 2.0 in 2020,^5^ there has been continued worldwide effort in testing and identification of EDCs, and this has resulted in additional large corpus of published literature. Therefore, in this study, we present an updated version, DEDuCTv3.0, that compiles information on 1043 potential EDCs from 3269 research articles published until January 2025.

Compared to the previous version, it was observed that the key sources for published literature such as World Health Organization (WHO) report,^23^ the Endocrine Disruption Exchange (TEDx; https://endocrinedisruption.org/) and EDCs Databank version 2015,^24^ were no longer updated or accessible. Therefore, in this study, the four-stage screening criteria was employed to curate EDCs and associated toxicity information from published literature available in PubMed (https://pubmed.ncbi.nlm.nih.gov/) (Supporting Information; Figure 1).

Briefly, a keyword search in PubMed on January 21, 2025, resulted in 29688 articles. Subsequently, 14295 new articles were first manually screened for EDC-relevant keywords, resulting in 9136 new articles at the end of stage 1 (Figure 1). In stage 2, 1795 new articles were identified to contain *in vivo* or *in vitro* EDC-relevant experimental evidence in humans or rodents (Figure 1). In stage 3, the chemical and associated toxicity information were manually curated from the full texts of these articles, and chemicals were further standardized using their two-dimensional (2D) structural identifiers from resources such as PubChem (https://pubchem.ncbi.nlm.nih.gov/) and Chemical Abstracts Service (CAS) Common Chemistry (https://commonchemistry.cas.org/) (Figure 1). Finally, in stage 4, chemicals were removed if they were either a natural hormone, mixture, or used as a therapeutic (Figure 1), and only chemicals comprising significant observed effects or endpoints related to endocrine disruption were retained, resulting in 1051 new articles containing information on 479 unique chemicals (Figure 1). Notably, this extensive manual screening procedure resulted in the identification of 251 novel EDCs, not compiled in previous versions of DEDuCT. Moreover, similar to the previous versions, the curated endocrine-mediated endpoints, and the corresponding dosage information have been standardized, and made publicly accessible via a dedicated webserver: https://cb.imsc.res.in/deduct/.

### 3.2. Construction of a toxicology knowledge graph for EDCs within DEDuCTv3.0

Toxicology knowledge graphs provide a framework to integrate and organize toxicological data from multiple sources, aiding in identification of underexplored relationships, and eventually enhancing our understanding of chemical-induced toxicities.^19^ In a previous study on reproductive toxicants, Evangelista *et al.*^19^ had constructed a knowledge graph linking chemicals, diseases, and genes from various resources, and utilized it to identify novel associations between these drugs and genes. Here, a knowledge graph was constructed using the chemical and endocrine-mediated endpoints data from DEDuCTv3.0, and subsequently utilized to explore novel endocrine-relevant gene interactions.

To construct the knowledge graph, several sources were relied upon to obtain semantic links between chemicals, endocrine-mediated endpoints, genes, phenotypes, diseases, Key Events (KE) and AOPs (Supporting Information; Figure 3). Briefly, the chemical-gene links were systematically curated from ToxCast invitrodb v4.3^12^ and CTD^14^ to retain human- or rodent-specific associations. Next, human- or rodent-relevant chemical-phenotype, chemical-disease, gene-phenotype, and gene-disease links were curated from CTD.^14^ Thereafter, the endocrine-mediated endpoint-phenotype and endocrine-mediated endpoint-disease links were manually inferred. Subsequently, the curated KE and AOP information from AOP-Wiki was utilized to identify links such as KE-AOP, chemical-AOP, phenotype-KE, disease-KE and endocrine-mediated endpoint-KE. The links between genes and KEs were obtained from annotated datasets provided by Saarimäki *et al.*^42^ Finally, to enable chemical similarity-based searches in the knowledge graph, the similarity between chemicals was computed as Tanimoto coefficient based on their ECFP4 fingerprints.^4,43^ Altogether, the constructed DEDuCT knowledge graph, referred to as ‘DEDuCT-KG’, comprised of 75106 unique nodes categorized under 7 node types, and 1065949 edges categorized under 31 edge types.

### 3.3. Exploration of potential EDCs across consumer products and chemical lists

The Chemical and Products Database v4.0 (CPDat), curated and maintained by the United States Environment Protection Agency (US EPA), contains information on the presence of chemicals in consumer products.^40^ Further, CPDat provides standardized product use categorization (PUC), that aids in characterizing the real-world chemical exposures.^40^ Therefore, CPDat was utilized to explore the exposure scenarios associated with potential EDCs within DEDuCTv3.0. In addition to the PUC data, the reported functional role of these chemicals was also obtained from CPDat.

In addition to consumer product exposure, the extent of regulatory coverage of these potential EDCs was explored across different chemical lists that were systematically compiled from inventories, guidelines and regulations from public resources (Table S6). Based on our earlier study,^5^ 54 chemical lists were compiled, among which 21 lists are classified as ‘Substances in Use’ (SIU), and 33 lists are classified as ‘Substances of Concern’ (SOC). Moreover, these lists have further been classified into ten broad categories based on their reported exposure scenario (Table S6). Next, the list of potential EDCs were compiled from DEDuCTv3.0, WHO report on EDCs,^23^ TEDx and EDCs Databank version 2015,^24^ and categorized as group I EDCs if they were present only in the SIU lists, group II EDCs if they were present in both SIU and SOC lists, or group III EDCs if they were present only in SOC lists. Finally, United States High Production Volume (USHPV) chemical list (Table S6) and the latest Organisation for Economic Co-operation and Development (OECD) High Production Volume (OECD HPV) chemical list (Table S6) were utilized to identify potential EDCs that are currently being produced in high volumes across the globe.

### 3.4. Construction of stressor-centric adverse outcome pathway networks for EDCs within DEDuCTv3.0

The Adverse Outcome Pathway (AOP) framework captures stressor-induced toxicity mechanisms, and organizes them across different levels of biological organization, terminating at an adverse outcome (AO) of regulatory relevance.^41^ The stressor-centric AOP network framework provides a holistic understanding of diverse AOPs associated with a stressor, eventually revealing the diversity in the stressor-induced toxicity mechanisms, and aiding in their risk assessment.^10,44^ In an earlier study, Ravichandran *et al.*^45^ explored the endocrine-relevant AOPs within AOP-Wiki, without revealing much on the mechanisms associated with individual EDCs within DEDuCT. Therefore, based on our earlier study,^10^ a systematic network-based filtration procedure was first leveraged to identify the complete and connected AOPs (Supporting Information).

In brief, the information within AOP-Wiki was first downloaded, and 385 complete and connected AOPs were systematically curated, and referred to as ‘curated AOPs’ in this study (Supporting Information; Tables S7-S9). Next, in addition to the curated endocrine-mediated endpoints within DEDuCTv3.0, human- and rodent-specific adverse effects were curated from ToxCast invitrodb v4.3,^12^ CTD (https://ctdbase.org),^14^ and NeurotoxKb (https://cb.imsc.res.in/neurotoxkb/),^46^ to obtain a holistic understanding of the toxicities associated with these EDCs. These endpoints were then systematically mapped with the biological key event (KE) information present within AOP-Wiki to obtain a stressor-centric AOP network linking 949 of 1043 EDCs in DEDuCTv3.0 to 381 curated AOPs (Supporting Information; Tables S10-S11). Metrics such as coverage score, i.e., a ratio of mapped events to the total events present within AOP, and level of relevance, i.e., a qualitative ranking to understand the kind of associated events, were utilized to characterize the identified EDC-AOP linkages in the constructed stressor-centric AOP network (Supporting Information; Table S11).^10^

### 3.5. Web interface and database management system

DEDuCT is an online resource accessible at: https://cb.imsc.res.in/deduct/. In this version update, the online resource has been enhanced with integration of all compile data generated in this study. In particular, the chemical section has been enhanced with more chemical-specific information pages on curated chemical-gene, chemical-phenotype and chemical-disease associations from CTD, curated ToxCast information from ToxCast,^12^ EDC-associated AOP data obtained from the constructed stressor-centric AOP network, and the consumer product presence information compiled from CPDat.^40^ The compiled data is stored in MariaDB (https://mariadb.org/), and retrieved using Structured Query Language (SQL). The web interface was created using HTML, CSS, Bootstrap4, jQuery (https://jquery.com/) and PHP (http://php.net/), and hosted on Apache webserver (https://httpd.apache.org/) running on Debian 9.4 operating system.

The constructed toxicology knowledge graph (DEDuCT-KG) is implemented using Neo4j (https://neo4j.com/). An open source web-based user interface, Knowledge-Graph-UI (KG-UI),^27^ was employed to create an interactive interface for DEDuCT-KG, which utilizes Cypher language (https://neo4j.com/docs/cypher-manual/current/introduction/) to query and retrieve the immediate neighbors and the shortest path between any two entities from the underlying knowledge graph (https://github.com/MaayanLab/Knowledge-Graph-UI/).

## 4. Conclusions

The Database of Endocrine Disrupting Chemicals and their Toxicity Profiles (DEDuCT), accessible at https://cb.imsc.res.in/deduct/, is an actively maintained and manually curated database on EDCs that has organized and structured data from scientific literature. In this study, we present an updated, enhanced and FAIR-compliant version, DEDuCTv3.0, which compiles 1043 chemicals from 3269 published studies and identifies 796 endocrine-mediated effects, covering published scientific literature until January 2025. In this update, EDC-specific toxicity data has been systematically collected from toxicology-relevant resources such as CTD, ToxCast, and AOP-Wiki, and made available on individual chemical pages, providing easy access to all chemical-related toxicity information. Additionally, the compiled data has been organized to build a large-scale EDC-specific toxicology knowledge graph, called DEDuCT-KG, and made accessible through an interactive interface. The construction of DEDuCT-KG has enabled exploration of potential obesogens within DEDuCT and supported the identification of their possible mechanisms of action. Furthermore, potential exposure sources were identified using the Chemical and Products Database (CPDatv4), showing exposure through everyday products such as cosmetics and home maintenance products, among others. Comparison with existing regulations also revealed chemicals that are currently overlooked. Overall, these analyses show that integrating diverse toxicological information has improved our understanding of the health effects associated with these potential EDCs and supports further research in this field.

While DEDuCT is a comprehensive manually curated resource on EDCs, it only focuses on compiling data from published scientific literature and does not include studies conducted by regulatory bodies or industries, which are typically published as chemical reports. Moreover, it primarily focuses on mammalian toxicities and does not address the ecotoxicity of such chemicals. It also does not cover the mixture effects of potential EDCs. Although the latest chemical lists were used for the regulatory analysis in this study, they might not fully capture all possible regulated chemicals.

Nonetheless, to the best of our knowledge, this study presents the first toxicology knowledge graph specifically developed for EDCs, termed DEDuCT-KG, enabling large-scale exploration of their adverse effects. DEDuCT-KG can facilitate the identification of mechanisms of action and uncover less-explored toxic effects associated with these chemicals. The new interactive user interface of DEDuCT-KG supports EDC research through specialized knowledge graph techniques. The inclusion of chemical structural similarity within DEDuCT-KG can aid in the construction of novel *in silico* tools that can be utilized for the characterization of EDCs. Moreover, the curated EDC-AOP associations will improve the understanding of chemical toxicities and help in designing targeted *in vitro* testing strategies for EDCs. Consumer product information from CPDat has allowed for the identification of exposure routes for several potential EDCs and highlighted primary sources of concern. In sum, this update of DEDuCT organizes the large amount of information on EDCs generated since the last version. It brings together diverse toxicology data into a central resource that can accelerate the research and regulation of EDCs.

## Data availability

The data associated with this study is contained in the article, or in the Supporting information files, or in the associated website: https://cb.imsc.res.in/deduct/, or in the associated GitHub repository: https://github.com/asamallab/DEDuCTv3.0.

## Acknowledgement

We would like to acknowledge U. Nagamalleswara Rao and G. Srinivasan for their guidance and help in setting up the Neo4j database and the knowledge graph user interface in the DEDuCT webserver. Areejit Samal and Nikhil Chivukula thank Janani Ravichandran for discussions. Areejit Samal would like to acknowledge funding from the Department of Atomic Energy (DAE), Government of India via Apex project to The Institute of Mathematical Sciences (IMSc) Chennai. The funders have no role in study design, data collection, data analysis, manuscript preparation or decision to publish.

## CRediT author contribution statement

**Nikhil Chivukula:** Conceptualization, Data Curation, Formal Analysis, Methodology, Software, Visualization, Writing; **Shrish Vashishth:** Data Curation; **Pavithra Kandasamy:** Data Curation; **Shreyes Rajan Madgaonkar**: Formal Analysis, Methodology, Software, Writing; **Areejit Samal:** Conceptualization, Supervision, Formal Analysis, Methodology, Writing.

## Supporting Figures Captions

**Figure S1.** (a) The distribution of the 796 endocrine-mediated endpoints in DEDuCTv3.0 into the seven systems-level perturbation categories. (b) The distribution of chemicals within DEDuCTv3.0 into the seven systems-level perturbation events based on their associated endocrine-mediated endpoint obtained from published literature. (c) A chronological analysis of the 3269 published articles supporting 1043 EDCs in DEDuCTv3.0. Note, 32 articles published in January of 2025 are not included. (d) Bar plot depicting number of new EDCs identified per year based on published literature compiled in DEDuCTv3.0. Note, 4 new EDCs from January 2025 are not included.

**Figure S2.** UpSet plot representation of unique overlaps between EDCs in DEDuCTv3.0 and other EDC-relevant databases such as the Endocrine Disruption Exchange (TEDx), EDCs Databank version 2015 (EDCsDB) and World Health Organization (WHO) report. (a) Comparison with all 1043 EDCs compiled in DEDuCTv3.0. (b) Comparison with the 251 new EDCs compiled in DEDuCTv3.0. These UpSet plots were generated using UpSetR package in R.

**Figure S3.** Network visualization of the chemical similarity network (CSN) constructed based on the Tanimoto coefficients computed for each pair of chemicals using ECFP4 fingerprints. In this figure, edges are filtered to represent only those with Tanimoto coefficient > 0.5, with the edge thickness proportionally scaled to the corresponding values. The chemicals compiled in DEDuCT 2.0 are represented in blue, while the 251 new chemicals curated in DEDuCTv3.0 are shown in red.

**Figure S4.** Network visualization of the target similarity network (CSN) constructed based on the Jaccard indices computed for each pair of chemicals using information on gene targets. In this figure, edges are filtered to represent only those with Jaccard index > 0.5, with the edge thickness proportionally scaled to the corresponding values. The 460 of 792 chemicals compiled in DEDuCT 2.0 are represented in blue, while the 119 of 251 new chemicals curated in DEDuCTv3.0 are shown in red.

**Figure S5.** The scatter plot of target similarity vs structure similarity between pairs of EDCs. Here, the structure similarity is given by Tanimoto coefficient, and target similarity is given by Jaccard Index. The plots are further divided based on different structural similarity cut-off values. The Pearson correlation coefficient (R) and the coefficient of determination (R2) values are denoted at the top-right of each graph. Further, the number of chemical pairs in each graph is denoted at the bottom right.

**Figure S6.** Distribution of 2049 unique potential EDCs from DEDuCTv3.0, TEDx, EDCs Databank, and WHO report across 54 chemical lists. (a) Venn diagram depicting chemicals categorized into groups I, II, and III. (b) Sunburst plots depicting distribution of potential EDCs across 10 categories of chemical lists. Within each category, the inner ring gives the number of potential EDCs categorized as groups I, II, or III, and the outer ring depicts the number of newly compiled chemicals within DEDuCTv3.0 present in each group.

**Figure S7.** Screenshot for the updated chemical information page in DEDuCTv3.0 webserver.

**Figure S8.** Screenshots from DEDuCTv3.0 webserver of the chemical information pages on: (a) chemical-gene interactions; (b) chemical-phenotype interactions; (c) chemical-disease associations.

**Figure S9.** Screenshot from DEDuCTv3.0 webserver of the chemical information page on associated ToxCast endpoints.

**Figure S10.** Screenshot from DEDuCTv3.0 webserver of the chemical information page on associated data from Chemical and Products Database (CPDat).

**Figure S11.** Screenshots from DEDuCTv3.0 webserver for the different search options. (a) Search based on chemical information such as names, synonyms, PubChem identifier, CASRN, DSSTox identifier, and DEDuCT identifier. (b) Search based on physicochemical properties of chemicals. (c) Search based on structural similarity.

## Supporting Tables Captions

**Table S1**: This table contains information on the 1043 EDCs compiled in DEDuCTv3.0. For each chemical, the table provides the chemical identifier, Chemical Abstracts Service Registry Number (CASRN), PubChem chemical identifier (CID), DSSTox identifier from EPA’s CompTox Chemicals Dashboard, chemical name, Bemis-Murcko scaffold represented as SMILES string, ClassyFire classifications (Kingdom, Superclass, Class, Subclass), evidence category based on the evidence compiled in DEDuCTv3.0 (I - evidence from in vivo human (IVH) studies; II - no evidence from IVH, but evidence from both in vitro human (IVTH), and in vivo rodent (IVR) studies; III - evidence only from IVR studies; IV - evidence only from IVTH studies), broad and sub-categories of the environmental presence information associated with the chemical, and the product use category (PUC) and functional use data (separated by ‘|’ symbol) obtained from Chemical and Products Database v4 (CPDat).

**Table S2**: This table contains the 796 curated endocrine-mediated effects in DEDuCTv3.0. For each endpoint, this table provides the corresponding systems-level perturbation(s) (separated by ‘;’ symbol), and the chemical identifier for the associated chemical(s) (separated by ‘|’ symbol).

**Table S3**: This table contains the No observed adverse effect level (NOAEL) and Low observed adverse effect level (LOAEL) values for EDCs based on test and effective dosage values reported in published experiments (accessible at https://cb.imsc.res.in/deduct/). Note that the supporting evidence for the EDCs in our resource has been compiled from three different types of published experiments or study types, namely, in vivo (IVH) or in vitro (IVTH) experiments in humans or in vivo (IVR) experiments in rodents. Importantly, NOAEL value for an EDC could not be determined from a published experiment if an endocrine-mediated endpoint or adverse effect is observed at every sampled test dosage, and in such a case, it is only possible to determine the LOAEL value for the EDC.

**Table S4**: This table contains information on the Chemical-Gene-Disease-Endpoint tetramer constructed for the ‘Lead to Obesity’ endocrine-mediated endpoint. For each tetramer, the table provides the chemical identifier, the NCBI gene identifier, the NCBI gene symbol, MeSH identifier of the disease, name fo the disease, and the endocrine-mediated endpoint from DEDuCT.

**Table S5**: This table provides information on the 18 identfied obesogens based on the knolwedge graph. For each chemical, the table provides the corresponding chemical identifier, Chemical Abstracts Service Registry Number (CASRN), name, number of associated genes identified from the tetramer, gene symbols of the associated genes (separated by ‘|’ symbol), and the literature source to support the chemical is classified as an obesogen.

**Table S6**: This table contains infromation on the chemical lists compiled in this study to perform the regulatory analysis. For each chemical list, the table provides the identifier, name, the source url from where the data was obtained, corresponding classification, and the type of the list.

**Table S7**: This table contains the curated list of 385 high confidence adverse outcome pathways from AOP-Wiki. For each AOP, the table provides the corresponding information on AOP identifier, AOP title, Handbook Version that was followed while construction of the AOP, Organisation for Economic Co-operation and Development (OECD) status, and taxonomic applicability of the AOP (separated by ‘|’ symbol). Additionally, for each AOP, the table provides the corresponding AOP identifier, computed fraction of KERs with ‘High’ evidence (i.e., F(High)), computed fraction of KERs with ‘Moderate’ evidence (i.e., F(Moderate)), computed fraction of KERs with ‘Low’ evidence (i.e., F(Low)), computed fraction of KERs with ‘Not Specified’ evidence (i.e., F(Not Specified)), computed cumulative weight of evidence (WoE).

**Table S8**: This table contains information on the 1228 Key Events (KEs) present in the curated list of 385 AOPs. For each KE, the table provides the corresponding information on KE identifier, KE title, level of biological organization, and associated AOP identifier(s) (separated by ‘|’ symbol).

**Table S9**: This table contains information on Key Event Relationships (KERs) present in each of the curated list of 385 AOPs. For each AOP, the table provides the AOP identifier, corresponding KER identifier, upstream KE identifier, downstream KE identifier, MIE(s) among upstream and downstream KEs (separated by ‘|’ symbol), AO(s) among upstream and downstream KEs (separated by ‘|’ symbol), adjacency of KER, weight of evidence of KER, and quantitative understanding of KER. Note that the KER identifiers starting with 10000 were manually assigned by the authors as the KER was mentioned in the AOP page but was not assigned an identifier in AOP-Wiki.

**Table 10:** This table contains information on 955 Key Events (KEs) from 381 high confidence AOPs that are associated with 949 EDCs. For each KE, the table provides corresponding information on KE identifier, chemical identifier (separated by ‘|’) for associated chemical(s), source(s) from which the associations are inferred (separated by ‘|’ symbol), AOP Identifier(s) from which KE mapping was inferred (separated by ‘|’ symbol), associated CTD disease(s) (separated by ‘|’ symbol), associated CTD phenotype(s) (separated by ‘|’ symbol), associated DEDuCT endpoint(s) (separated by ‘|’ symbol), associated NeurotoxKb endpoint(s) (separated by ‘|’ symbol), and associated ToxCast assay endpoint(s) (represented as ‘assay_name:response’)(separated by ‘|’ symbol).

**Table 11:** This table provides edge list for the complete stressor-AOP network for 949 EDCs within DEDuCTv3.0. For each edge in the network, the table provides stressor chemical identifier, computed coverage score, level of relevance, AOP identifier, and the AO classification of the corresponding AOP.

