## Supplementary material for "DEDuCT 3.0: An enhanced and expanded FAIR-compliant resource and toxicology knowledge graph for endocrine disrupting chemicals": Figure S

### for

### Supporting Text

#### 1. Workflow for the compilation of Endocrine Disrupting Chemicals (EDCs)

The Database of Endocrine Disrupting Chemicals and their Toxicity profiles (DEDuCT) is a comprehensive knowledgebase that systematically compiles information on potential Endocrine Disrupting Chemicals (EDCs) from published literature.[1,2] The latest version, DEDuCT version 2.0 (released on October 2, 2020) compiled information on 792 potential EDCs with supporting evidence from 2218 articles.[2] Since its release, global efforts in testing and identification of EDCs have generated an additional large corpus of published literature. Therefore, in this study, we present an updated version, DEDuCT version 3.0 (DEDuCTv3.0), that compiles information on 1043 potential EDCs from 3269 research articles published until January 2025. To achieve this, a systematic four-staged workflow based on extensive manual curation was utilized to curate chemicals and their associated endocrine-mediated effects in humans or rodents from published literature (Figure 1).[1] This four-staged workflow is summarized in the following sections.

##### 1.1. Stage 1: Mining EDC-relevant literature

The first stage involves the mining of EDC-relevant published literature from different sources. In comparison to the previous study, it was observed that the sources for published literature, such as the World Health Organization (WHO) report,[3] the Endocrine Disruption Exchange (TEDx, https://endocrinedisruption.org/) and EDCs Databank version 2015,[4] were no longer updated or accessible. Therefore, only PubMed (https://pubmed.ncbi.nlm.nih.gov/) was queried with the following query:

"EDCs" OR "EDC" OR ("endocrine" AND "disrupt") OR ("disrupt" AND "endocrine") OR "endocrine disruptors" OR "endocrine-disruptors" OR "endocrine disruptor" OR "endocrine-disruptor" OR "endocrine disrupters" OR "endocrine-disrupters" OR "endocrine disruption" OR "endocrine-disruption" OR "endocrine disruptive" OR "endocrine-disruptive" OR "endocrine disrupting" OR "endocrine-disrupting" OR "endocrine disrupter"

This search, last conducted on January 21, 2025, resulted in a list of 29688 published literature, among which 14295 articles were not screened in the previous two versions of DEDuCT. Thereafter, the title and abstracts of these 14295 new articles were manually screened for keywords relevant to EDCs, resulting in 9136 new articles at the end of stage 1 (Figure 1).

#### 1.2. Stage 2: Literature filter based on study type and test organism

In stage 2 of the workflow, 9136 new articles filtered from stage 1 were screened to identify studies based on *in vivo* or *in vitro* experiments in humans or rodents. During this stage, the studies comprising receptor-based binding assays or *in silico* methods to infer endocrine disrupting potential were filtered out as the evidence is insufficient to determine if the chemical can lead to adverse effects based on environmental exposures. Through this extensive screening process, 1795 new research articles relevant to EDC-specific experiments in humans or rodents were obtained (Figure 1).

#### 1.3. Stage 3: Compilation of tested chemicals from the filtered research articles

In stage 3 of the workflow, the full text of the filtered 1795 articles were first obtained for comprehensive screening of tested chemicals. It was observed that full texts were not available for nine articles and were therefore excluded from further analysis. Chemical data was curated from the remaining 1786 articles, and were mapped to their two-dimensional (2D) structure identifiers using standard chemical databases such as PubChem (https://pubchem.ncbi.nlm.nih.gov/) and Chemical Abstracts Service (CAS) (https://commonchemistry.cas.org/). This resulted in the identification of 637 unique chemicals that have been tested for endocrine disruption in humans or rodents in at least one research article among the 1786 screened articles obtained after stage 2 (Figure 1).

#### 1.4. Stage 4: Identification of EDCs with supporting evidence on systems-level endocrine-mediated perturbations

In stage 4 of the workflow, the tested chemicals were assessed for the presence of significant observed endocrine-associated adverse effects in the published literature. First, the chemicals were discarded if they were tested as part of a mixture, or were natural hormones, or were tested for therapeutic usage in the associated literature. Next, studies were discarded if they solely relied on *in vitro* rodent experiments. Thereafter, the observed adverse effects or endpoints were first checked for significance within the associated literature and then checked for endocrine disruption relevance, such as changes to morphology, physiology, reproduction, growth, development, and lifespan. Finally, a chemical was considered as a potential EDC if it contained at least one significant supporting evidence for endocrine-relevant effects from associated literature. Through this extensive manual curation, 479 chemicals were identified with strong supporting evidence for endocrine disruption from 1051 published articles.

Next, the reported adverse effects associated with these 479 chemicals were manually standardized based on the vocabulary established in earlier versions of DEDuCT,[1,2] resulting in the compilation of 573 endocrine-mediated endpoints. Among these 573 endpoints, 187 were not present in earlier versions, and were therefore manually classified into the 7 systems-level perturbations based on the major biological processes controlled by the human endocrine system, namely, Endocrine-mediated cancer (CT), Reproductive (RT), Developmental (DT), Metabolic (MT), Immunological (IT), Neurological (NT), Hepatic (HT) endocrine-mediated perturbations. These unified terms can serve as standard biological vocabulary describing the toxicity profiles of EDCs.[1] In addition to the compilation of endocrine-mediated endpoints, the corresponding dosage ranges were also compiled from the supporting literature for these 479 potential EDCs. The dosage units were uniformized, and the dosages were recorded as test and effective dosage for each EDC to enable dose-response measurements.

Finally, the compiled data was integrated with DEDuCT version 2.0, resulting in a list of 1043 unique potential EDCs, comprising 796 unique endocrine-mediated endpoints systematically compiled from 3269 published literature.

A chronological analysis of the 3269 articles revealed that nearly half the 3269 articles compiled in DEDuCTv3.0 were published between 2017 and January of 2025 (Figure S1c). Notably, it was observed that the reliance on *in vitro* human experiments is gradually increasing over the years, suggesting the reliance on non-animal approaches towards EDC research (Figure S1c). In addition, the literature data was utilized to understand the identification of new EDCs over the years. Here, the supporting evidence for each EDC was sorted chronologically, and the year of identification was defined as the date of the earliest published literature. Through this analysis, it was observed that the number of new EDCs identified has increased over the past few decades, aligning with the growth in research and interest regarding endocrine disruptors (Figure S1d).

### 2. Additional information on EDCs in DEDuCTv3.0

In addition to experimental evidence, DEDuCTv3.0 also compiles diverse information for the 1043 potential EDCs including two-dimensional (2D) and three-dimensional (3D) chemical structures, physicochemical properties, predicted ADMET properties, molecular descriptors, associated gene, phenotype and disease information curated from Comparative Toxicogenomics Database (CTD)[3] (https://ctdbase.org), active readouts from ToxCast invitrodb v4.3, associated Adverse Outcome Pathways (AOPs) from AOP-Wiki (Table S11), consumer products presence information from CPDat v4, and presence in various regulation lists and in human biospecimens.

The potential EDCs are classified based on their environmental sources into 7 broad categories and 48 sub-categories (Table S1). Further, ClassyFire[6] was utilized to hierarchically classify the EDCs based on their structure (Table S1). In case ClassyFire[6] failed to classify the chemical, the structural definitions were relied upon to obtain the Kingdom of such chemicals.[6] Moreover, the 1043 potential EDCs were classified into 4 categories (I-IV) based on the type of supporting evidence for endocrine disruption in published experiments specific to humans or rodents (Table S1). The compiled information in DEDuCT can be downloaded in flat file format from the associated webserver at: https://cb.imsc.res.in/deduct/download.

In sum, we provide an expanded list of 1043 potential EDCs in DEDuCTv3.0, which could assist academia, industry, and regulatory agencies to develop safer consumer products.

**3. Exploration of chemical space within DEDuCTv3.0**

The chemical space of 1043 EDCs within DEDuCTv3.0 was explored using a chemical similarity network (CSN) and a target similarity network (TSN). To construct the CSN, 1043 EDCs were used as nodes, and the edges denoted structural similarity between corresponding nodes. Based on the previous study, the chemical similarities were computed between chemicals as the Tanimoto coefficient based on their ECFP4 fingerprints.[7,8] Thereafter, the chemical pairs having Tanimoto coefficients > 0.5 were considered as being structurally similar, and were used as edge weights in the network (Figure S3).

To construct TSN, the chemical-gene interaction information was curated from ToxCast invitrodb v4.3,[9] as explained in section 7.4 titled 'Identification of KEs using ToxCast'. This resulted in the identification of 579 of 1043 EDCs targeting 400 human or rodent relevant genes. Based on the previous study,[7] the similarities between chemicals were computed as the Jaccard index[10] of the corresponding target gene sets. Thereafter, the chemical pairs having a Jaccard index > 0.5 were considered as being target-wise similar, and were used as edge weights in the network (Figure S4).

Based on the previous study,[7] the correlation between structural similarity and target similarity was further probed for 579 of 1043 chemicals with target gene information from

ToxCast (Figure S11). The structural similarity based on the Tanimoto coefficient was plotted on the y-axis, and the target similarity based on the Jaccard Index was plotted on the x-axis. Thereafter, Person correlation coefficient was computed to check for correlation between structural and target similarities.

**4. Compilation of AOPs within AOP-Wiki**

Adverse Outcome Pathway (AOP) is a toxicological knowledge framework that captures biological processes underlying stressor-induced toxicity.[11] In this framework, toxicity-related biological events, referred to as Key Events (KEs), are organized sequentially, beginning with the interaction of the stressor with a biological target, known as the Molecular Initiating Event (MIE), and culminating in an Adverse Outcome (AO).[11–14] The causal, directional relationships between these events are termed Key Event Relationships (KERs).[11–14] AOPs are stressor-agnostic and context-specific, meaning each AOP is tailored to a particular biological scenario.[11–14] Developed globally, these AOPs are deposited in the AOP-Wiki (https://aopwiki.org), which is the largest, publicly accessible repository hosted by the Society for the Advancement of Adverse Outcome Pathways (SAAOP). AOP-Wiki hosts several AOPs, where each AOP is documented in the form of KEs (including MIE and AO) and KERs, all of which are supported by scientific evidence (https://aopwiki.org/handbooks/5). Therefore, in this study, AOP-Wiki was relied upon to obtain the latest available AOPs.

First, the XML file (released April 1, 2025) was downloaded from the 'Project Downloads' page in AOP-Wiki to obtain the latest information on AOPs. Then, the data was parsed to extract information associated with AOPs like AOP identifier, AOP title, associated KEs (including MIEs and AOs) and KERs, linked stressors, handbook version followed for development of AOP, status according to OECD, and biological applicability information such as taxonomy, sex and life-stage of the organism, and their corresponding weight of evidence, using an in-house Python script. Additionally, for each KE, information like KE title, KE identifier, level of biological organization, action name, object name, object identifiers, and process name, and information associated with KERs like upstream/downstream KEs, evidence for biological plausibility of KER, adjacency, and the extent of quantitative understanding of KER, were extracted.

**5. Curating complete and connected AOPs within AOP-Wiki**

AOPs are considered living documents as they are collaboratively developed globally, and are continuously updated based on new scientific evidence (https://aopwiki.org/handbooks/5). Consequently, many AOPs may remain incomplete or lack sufficient information for further analysis. Therefore, based on our previous studies,[15–17] each AOP was systematically checked to filter high quality and complete AOPs. First, AOPs were manually checked and those that comprised KEs with title as 'unknown', or lacked any KEs or KERs, were removed. Next, the NetworkX library[18] was employed in Python to check for disconnected AOPs, and those that contained disconnected components were manually inspected and updated before filtration of AOPs with disconnected components. Finally, the presence of MIE, AO, and a directed path between them was checked, and those that lacked any were filtered out. Through this combined manual and computational effort, 385 complete, connected and high quality AOPs were obtained from AOP-Wiki, which were then designated as 'curated AOPs' (Table S7). These 385 high confidence AOPs comprised 1228 unique KEs (Table S8) and 1966 unique KERs (Table S9).

### 6. Computation of cumulative weight of evidence (WoE) of AOPs

AOP-Wiki provides a weight of evidence (WoE) information for each KER in an AOP based on its biological plausibility (Table S9). The WoE is a qualitative score represented in terms of 'High', 'Moderate', 'Low', and 'Not Specified'. Following Ravichandran *et al.*,[19] the fraction of KERs within an AOP with 'High' WoE [represented as F(High)], 'Moderate' WoE [represented as F(Moderate)], 'Low' WoE [represented as F(Low)] and 'Not Specified' WoE [represented as F(Not Specified)], were computed, and subsequently, a cumulative WoE was assigned to the AOPs based on the following criteria:

i. If $F(High) \geq 0.5$, the cumulative WoE of AOP is 'High'
ii. If $F(High) < 0.5$, but $(F(High) + F(Moderate)) \geq 0.5$, the cumulative WoE of AOP is 'Moderate'
iii. If $(F(High) + F(Moderate)) < 0.5$, but $(F(High) + F(Moderate) + F(Low)) \geq 0.5$, the cumulative WoE of AOP is 'Low'
iv. If none of the above criteria are satisfied, the cumulative WoE of AOP is 'Not Specified'

Table S7 provides the cumulative WoE for the curated AOPs.

### 7. Identification of KEs associated with EDCs

Based on our previous studies,[15–17] different toxicogenomics and biological endpoint information related to EDCs was obtained from AOP-Wiki (https://aopwiki.org), Comparative Toxicogenomics Database (CTD)[5] (https://ctdbase.org), DEDuCT[1,2] (https://cb.imsc.res.in/deduct/), NeurotoxKb[20] (https://cb.imsc.res.in/neurotoxkb/) and ToxCast invitrodb v4.3.[21] The obtained data was then systematically integrated to identify KEs within AOP-Wiki that are linked to EDCs.

#### 7.1. Identification of KEs using AOP-Wiki

AOP-Wiki catalogs information on prototypical stressors for each AOP, based on well-documented associations with these stressors (https://aopwiki.org/handbooks/5). In this study, the information on prototypical stressors associated with AOPs was extracted from the flat download file using an in-house Python script. Thereafter, 100 EDCs were identified as prototypical stressors for 93 AOPs. Subsequently, the 361 KEs within these AOPs were considered as associated with these 100 EDCs (Table S10).

#### 7.2. Identification of KEs using CTD

The Comparative Toxicogenomics Database (CTD) is a comprehensive public resource that links environmental chemicals (C), genes (G), phenotypes (P), and diseases (D) to advance understanding of their effects on health.[5] This resource facilitates the construction of CGPD-tetramers, which help identify confident associations between chemicals, phenotypes, and diseases, enabling their mapping to KEs within AOP-Wiki. Based on our previous work,[15–17] high confidence CGPD-tetramers associated with EDCs were retrieved and subsequently leveraged to identify KEs within AOP-Wiki.

CTD provides a chemical dictionary linking standard PubChem identifiers and CASRN to chemical IDs within CTD. Initially, a list of 804 unique CTD chemical identifiers was obtained based on overlaps with data compiled in DEDuCTv3.0. Upon closer inspection, it was observed that some of the associations provided by CTD are incorrect. For instance, the EDC Benzo(a)pyrene (DED000576) is linked to two chemicals IDs within CTD based on its PubChem identifier (CID:2336). Upon closer inspection, it was observed that one of the chemical ids, MESH:D001580 corresponds to the class of benzopyrenes, while MESH:D001564 corresponds to Benzo(a)pyrene. Therefore, all the associations were manually screened to remove non-specific information within CTD, resulting in the identification of 783 chemical ids within CTD associated with 764 EDCs within DEDuCTv3.0.

Next, the CTD November 2025 release was accessed and the CGPD-tetramers were constructed, wherein the following were considered: (i) chemical-gene and chemical-phenotype associations that are relevant to humans and rodents, and are supported with literature evidence; (ii) chemical-disease and gene-disease associations with 'marker/mechanism' evidence, and are supported with literature evidence; (iii) gene-phenotype associations with GO annotations based on only the experimental results in human or rodent-based studies (https://geneontology.org/docs/guide-go-evidence-codes/). This process resulted in a list of 208809 CGPD-tetramers comprising 364 EDCs, 3311 genes, 1747 phenotypes, and 858 diseases. Next, the immediate neighboring GO terms for the CGPD-tetramer phenotype GO terms were generated using the GOSim package[22] in the R programming language. These GO terms were then overlapped with the process identifiers of KEs in AOP-Wiki, and manually screened to identify 310 KEs linked to 279 phenotypes across 347 EDCs (Table S10). Additionally, the disease terms were manually mapped to identify 256 KEs associated with 548 diseases across 358 EDCs (Table S10).

#### 7.3. Identification of KEs using DEDuCTv3.0 endocrine-mediated endpoints and NeurotoxKb neurotoxicant endpoints

Based on previous studies,[15–17] the 796 endocrine-mediated endpoints associated with the 1043 potential EDCs within DEDuCTv3.0, were first screened to identify relevant KEs within AOP-Wiki. In particular, the endpoints and the KE titles were manually inspected, resulting in the identification of 276 KEs associated with 361 endocrine-mediated endpoints, across 927 EDCs (Table S10).

NeurotoxKb[20] (https://cb.imsc.res.in/neurotoxkb/) is a manually curated resource focusing on mammalian neurotoxicity endpoints associated with environmental chemicals, compiled from published literature. In this study, the neurotoxic endpoints corresponding to EDCs within NeurotoxKb were extracted, and subsequently considered to identify relevant KEs within AOP-Wiki. The neurotoxic endpoints and KE titles in AOP-Wiki were manually inspected, to identify 64 KEs associated with 55 neurotoxic endpoints across 123 EDCs (Table S10).

#### 7.4. Identification of KEs using ToxCast

ToxCast is a program by the United States Environmental Protection Agency (US EPA) designed to enhance chemical toxicity predictions through *in vitro* high-throughput screening of various environmental chemicals.[21] These *in vitro* approaches provide crucial information on the associated biological processes and gene alterations triggered by stressor interactions, which can aid in the identification of potential MIEs related to active chemicals.[15–17,23–25] In this study, ToxCast invitrodb v4.3[9,26] was utilized to identify KEs associated with EDCs.

First, the chemicals and their corresponding assay information were extracted from the 'mc5-6_winning_model_fits-flags_invitrodbv4_3_AUG2024.csv' file, and active assay endpoints for each chemical (defined as 'hitc' ≥ 0.9) were identified.[26] Next, the 'top' value of the corresponding winning model from the 'mc4_all_model_fits_invitrodbv4_3_AUG2024.Rdata' file was utilized to determine whether these active chemicals exhibited an 'activatory' or 'inhibitory' effect.[26]

To ensure that these active endpoints are not due to non-specific activation of reported genes (referred to as 'cytotoxicity-associated burst' phenomenon),[27] the following Z-score statistic proposed by Judson *et al.*[27] was applied:

$$Z(chemical, assay) = \frac{-logAC_{50}(chemical,assay) - median[-logAC_{50}(chemical,cytotox)]}{\text{global cytotoxicity MAD}}$$

Here, '$\log AC_{50}$(chemical,assay)' is the logarithm of the $AC_{50}$ value of the chemical in the assay, '$\log AC_{50}$(chemical,cytotox)' is the logarithm of the $AC_{50}$ value of the chemical in the corresponding cytotoxicity assay, and the 'global cytotoxicity MAD' is the median of the median average deviations (MAD) of the $\log AC_{50}$(chemical,cytotox) distributions across all chemicals. Based on this definition, Z-scores ranging between +3 and -3 are considered indicative of cytotoxicity-associated bursts.[27]

In this study, the global cytotoxicity MAD and $\log AC_{50}$(chemical,cytotox), given by the column titled 'cytotox_median_log', were retrieved from the 'cytotox_invitrodb_v4_3_AUG2024.xlsx' file to identify the cytotoxicity-associated bursts associated with EDCs.[26] The assays with Z-scores between +3 and -3 were discarded, and the resulting assays were subsequently filtered to obtain human- or rodent-specific endpoints, and then mapped to KEs within AOP-Wiki.

The ToxCast assay endpoints and the KE titles from the AOP-Wiki were manually inspected to identify relevant KEs. This manual curation led to the identification of 160 KEs associated with 318 assay endpoints across 523 EDCs (Table S10).

Altogether, 955 KEs were identified to be associated with 984 EDCs through the integration of heterogeneous information on toxicogenomic and biological endpoints from five exposome-relevant resources, namely, AOP-Wiki, CTD, DEDuCT, NeurotoxKb, and ToxCast (Table S10).

### 8. Construction of stressor-AOP network

Stressor-AOP network provides a broader perspective on the impacts of stressors across diverse biological processes by linking the stressors to AOPs.[16,24] To better understand the perturbances caused by EDCs, a stressor-AOP network was constructed, linking EDCs to different AOPs within AOP-Wiki. In order to obtain high confidence associations between the chemicals and AOPs, only the 385 curated AOPs were relied upon (Table S7).

First, the EDCs were linked to any AOP if they shared at least one common KE. Thereafter, these links were characterized based on two criteria namely, the coverage score and the level of relevance. The coverage score of a stressor-AOP link is defined as the ratio of the number of KEs within that AOP associated with the stressor to the total number of KEs within that AOP.[15,28] This score is a real-valued number between 0 and 1, and is denoted as the edge weight of the linkage between a stressor and an AOP in the constructed stressor-AOP network. The level of relevance is a qualitative score used to identify the relevance of stressor-AOP association within the stressor-AOP network,[16] and this score is denoted as an attribute of the edge in the constructed stressor-AOP network. Level of relevance is a five-level criterion defined as follows:

- *Level 1*: The stressor is associated with at least one KE within an AOP, where the KE is neither MIE nor AO within that AOP
- *Level 2*: The stressor is associated with at least one AO within an AOP, but not associated with any MIE within that AOP
- *Level 3*: The stressor is associated with at least one MIE within an AOP, but not associated with any AO within that AOP
- *Level 4*: The stressor is associated with at least one MIE and one AO within an AOP
- *Level 5*: The stressor is associated with at least one MIE and one AO within an AOP, and there exists a directed path between the associated MIE and AO

Table S11 contains all the data on the stressor-AOP network constructed for EDCs in DEDuCTv3.0, including the coverage score and level of relevance for each of the stressor-AOP links.

### 9. Construction of a toxicology knowledge graph for EDCs in DEDuCTv3.0

In this study, a toxicology knowledge graph was constructed by linking the chemicals (EDCs) and endocrine-mediated endpoints within DEDuCTv3.0 with gene, phenotype, disease, biological key events (KEs), and adverse outcome pathways (AOPs) from toxicology-relevant resources such as ToxCast, CTD, and AOP-Wiki (Figure 4). We refer to this knowledge graph for EDCs as 'DEDuCT-KG', and the following subsections summarize the steps involved in curation of links and nodes within the knowledge graph.

#### 9.1. Curation of links between chemicals and endocrine-mediated endpoints

The endocrine-mediated endpoints associated with chemicals have been systematically curated from published literature in DEDuCTv3.0, and therefore can be considered as high confidence links. All these chemical-endpoint links have been assigned the edge type 'causes' to better explain these links in the knowledge graph. Through this process, 10532 links were identified, that connected 1043 EDCs with 796 endocrine-mediated endpoints in DEDuCT-KG.

#### 9.2. Curation of links between chemicals and genes from CTD and ToxCast

CTD provides curated chemical-gene interactions obtained from published literature, where it compiles additional information on the gene form of the interacting gene, corresponding organism name, and taxonomic identifier, chemical-gene interaction actions, and associated PMID of the published literature. First, the NCBI taxonomy browser (https://www.ncbi.nlm.nih.gov/taxonomy) was searched with the corresponding organism identifiers to obtain information on the taxonomic classification such as the domain, clade, kingdom, phylum, and order, among others. To obtain human- or rodent-relevant gene information, only species such as *Homo sapiens* (organism id: 9606) and those belonging to the order 'Rodentia' were selected. This resulted in the identification of 28 relevant species. Next, genes were filtered based on their gene form information to obtain those entries that were annotated as gene, mRNA, or protein. Further, genes were filtered out based on the associated interaction actions such as 'abundance', 'synthesis', 'response to substance', 'reaction', 'cotreatment', 'transport', 'localization', 'degradation', 'export', 'import', 'metabolic processing', 'mutagenesis', 'secretion', 'uptake', 'stability', and only those with terms such as 'expression' and 'activity' were retained. These interaction actions further included the kind of actions, such as 'increases', 'decreases', or 'affects'. For some of the chemical-gene links, it was observed that there were different curated interaction actions across different studies. Therefore, to mitigate such discrepancies, the following standardization criteria was followed: (i) if the interaction action included terms such as both 'increases'/'decreases' and 'affects', then it was assigned to 'increases'/'decreases' respectively; and (ii) in case both 'increases' and 'decreases' were present, then it was assigned as 'affects'. Through this systematic standardization process, 509996 links were identified that connected 682 chemicals to 29560 genes. The standardized interaction actions were assigned as the edge types within the knowledge graph.

Chemical-gene interactions were also obtained from human- or rodent-relevant data curated from ToxCast invitrodb v4.3. Here, the endpoints where the response was characterized as either 'activatory' or 'gain' were assigned as 'activates', and those characterized as 'inhibitory' or 'loss' were assigned as 'inhibits'. Through this process, 13205 links were identified, that connected 579 chemicals to 400 genes. The standardized response information was assigned as edge types within the knowledge graph.

#### 9.3. Curation of links between chemicals and phenotypes or diseases from CTD

CTD provides curated chemical-phenotype interactions obtained from published literature, where it compiles additional information on the corresponding organism name and taxonomic identifier, chemical-phenotype interaction actions, and associated PMID of the published literature. Similar to chemical-gene links, these interactions were first filtered to obtain human- or rodent-relevant information. Subsequently, the discrepancies in chemical-phenotype interactions were resolved, resulting in the identification of 28561 links that connected 599 chemicals to 4431 phenotypes. The standardized interaction actions were assigned as the edge types within the knowledge graph.

CTD also provides curated chemical-disease associations, where it additionally annotates the association as either 'marker/mechanism' or 'therapeutic' based on the evidence from corresponding published literature. According to the CTD terminology, 'marker/mechanism' corresponds to evidence suggesting that the chemical exposure correlates with the disease or may play a role in the etiology of the disease, while 'therapeutic' corresponds to evidence suggesting the therapeutic potential of chemical in a disease (https://ctdbase.org/help/glossary.jsp). Therefore, only associations marked as 'marker/mechanism' were obtained, resulting in the identification of 16219 links connecting 586 chemicals to 1805 diseases. The edge type was assigned as 'causes_or_correlated_with' to better explain the chemical-disease associations in the knowledge graph.

#### 9.4. Curation of links between genes and disease within CTD

CTD provides curated gene-disease associations, where it additionally annotates the association as either ‘marker/mechanism’ or ‘therapeutic’ based on the evidence from corresponding published literature. According to CTD terminology, ‘marker/mechanism’ corresponds to evidence suggesting that the gene may be a biomarker for the disease or may play a role in the etiology of the disease, while ‘therapeutic’ corresponds to evidence suggesting the gene may be a potential therapeutic target in the treatment of the disease (https://ctdbase.org/help/glossary.jsp). Therefore, only associations marked as ‘marker/mechanism’ were obtained, resulting in the identification of 27598 links connecting 7772 genes to 2466 diseases. The edge type was assigned as ‘associated_with’ to better explain the gene-disease associations in the knowledge graph.

#### 9.5. Curation of links between genes and phenotypes from Gene Ontology

Gene Ontology (GO) provides annotated gene-phenotype associations, including the taxonomy identifier of the organism in which the association was identified, the kind of evidence and a qualifier term to explain the association between the gene and phenotype. First, the GO annotations were obtained from the NCBI gene resource (https://ftp.ncbi.nih.gov/gene) (last accessed on November 26, 2025), and only gene-phenotype associations with experimental results were retained (https://geneontology.org/docs/guide-go-evidence-codes/). Next, the links were filtered to retain only human- or rodent-relevant associations based on taxonomic information. It was observed that each link was annotated with a qualifier term that defines the gene-phenotype association. The links were then filtered to remove qualifier terms that included ‘NOT’, resulting in the identification of 304398 links connecting 36463 genes to 18311 phenotypes. The qualifier terms were then assigned as the edge types within the knowledge graph.

#### 9.6. Curation of links between endocrine-mediated endpoints and phenotypes or diseases

In DEDuCTv3.0, endocrine-mediated endpoints were manually curated based on the biological endpoints reported in the supporting information. Mapping these endpoints to phenotypes and disease terms enables making the data more FAIR-compliant.[29] For this purpose, the definitions of the endpoints, phenotypes, and disease terms were manually checked and mapped based on underlying biological similarities. Through this process, 581 links were identified that connected 394 endocrine-mediated endpoints to 248 phenotypes and 78 diseases. These links have been assigned the edge type 'related_to' to better explain these links in the knowledge graph.

#### 9.7. Curation of links between chemicals, genes, phenotypes, diseases, endocrine-mediated endpoints, and key event information within AOP-Wiki

Integrating diverse toxicological data with information contained within the AOP-Wiki enables the exploration of diverse toxicity mechanisms associated with chemicals.[15–17,30] Furthermore, integrating these data within a knowledge graph enables the exploration of previously missed associations and may potentially aid in elucidating the chemical-associated toxicity mechanisms that were otherwise less understood. Based on the procedure explained in the section 7 of supporting information titled 'Identification of KEs associated with EDCs', all the endocrine-mediated endpoints, phenotypes, and diseases were systematically mapped to KE information within AOP-Wiki. This resulted in the identification of 2334 links connecting 411 phenotypes, 733 diseases, 320 endpoints, with 632 KEs within AOP-Wiki. These links have been assigned the edge type 'related_to' to better explain these links in the knowledge graph. Next, the curated prototypical stressor information from AOP-Wiki were utilized to identify 175 links connecting 100 chemicals to 93 AOPs. These links have been assigned the edge type as 'prototypical_stressor_of' to better explain these links in the knowledge graph.

Further, the gene-KE associations were obtained from data curated by Saarimäki *et al.*[31] (https://doi.org/10.5281/zenodo.7980953). In this dataset, the provided Ensembl gene IDs were mapped to Entrez gene identifiers using the biomart online tool (https://asia.ensembl.org/info/data/biomart/index.html?) (last accessed on November 26, 2025), resulting in the identification of 148171 links connecting 969 KEs and 14298 genes. These links have been assigned the edge type 'mapped_to' to better explain these links in the knowledge graph.

#### 9.8. Curation of links between chemicals based on structural similarity

To enable chemical similarity-based search across DEDuCT-KG, the chemical similarities were computed between EDCs as Tanimoto coefficient based on their ECFP4 fingerprints.[7,8] Subsequently, the chemical pairs having Tanimoto coefficients > 0.5 were considered as having structural similarity. This resulted in the identification of 1609 links connecting 634 chemicals within DEDuCTv3.0. These links have been assigned the edge type 'structurally_similar_to' to better explain these links in the knowledge graph.

(13) Villeneuve, D. L.; Crump, D.; Garcia-Reyero, N.; Hecker, M.; Hutchinson, T. H.; LaLone, C. A.; Landesmann, B.; Lettieri, T.; Munn, S.; Nepelska, M.; Ottinger, M. A.;

Vergauwen, L.; Whelan, M. Adverse Outcome Pathway Development II: Best Practices. *Toxicological Sciences* **2014**, *142* (2), 321–330. https://doi.org/10.1093/toxsci/kfu200.
(14) OECD. Revised Guidance Document on Developing and Assessing Adverse Outcome Pathways, Series on Testing and Assessment No. 184. *OECD Publishing, Paris* **2017**.
(15) Sahoo, A. K.; Chivukula, N.; Ramesh, K.; Singha, J.; Marigoudar, S. R.; Sharma, K. V.; Samal, A. An Integrative Data-Centric Approach to Derivation and Characterization of an Adverse Outcome Pathway Network for Cadmium-Induced Toxicity. *Science of The Total Environment* **2024**, *920*, 170968. https://doi.org/10.1016/j.scitotenv.2024.170968.
(16) Sahoo, A. K.; Chivukula, N.; Madgaonkar, S. R.; Ramesh, K.; Marigoudar, S. R.; Sharma, K. V.; Samal, A. Leveraging Integrative Toxicogenomic Approach towards Development of Stressor-Centric Adverse Outcome Pathway Networks for Plastic Additives. *Archives of Toxicology* **2024**, *98* (10), 3299–3321. https://doi.org/10.1007/s00204-024-03825-z.
(17) Sahoo, A. K.; Madgaonkar, S. R.; Chivukula, N.; Karthikeyan, P.; Ramesh, K.; Marigoudar, S. R.; Sharma, K. V.; Samal, A. Network-Based Investigation of Petroleum Hydrocarbons-Induced Ecotoxicological Effects and Their Risk Assessment. *Environment International* **2024**, *194*, 109163. https://doi.org/10.1016/j.envint.2024.109163.
(18) Hagberg, A. A.; Schult, D. A.; Swart, P. J. Exploring Network Structure, Dynamics, and Function Using NetworkX. In *Proceedings of the 7th Python in Science Conference (SciPy 2008)*; Varoquaux, G., Vaught, T., Millman, J., Eds.; 2008; pp 11–15.
(19) Ravichandran, J.; Karthikeyan, B. S.; Samal, A. Investigation of a Derived Adverse Outcome Pathway (AOP) Network for Endocrine-Mediated Perturbations. *Science of the Total Environment* **2022**, *826*, 154112. https://doi.org/10.1016/j.scitotenv.2022.154112.
(20) Ravichandran, J.; Karthikeyan, B. S.; Singla, P.; Aparna, S. R.; Samal, A. NeurotoxKb 1.0: Compilation, Curation and Exploration of a Knowledgebase of Environmental Neurotoxicants Specific to Mammals. *Chemosphere* **2021**, *278*, 130387. https://doi.org/10.1016/j.chemosphere.2021.130387.
(21) Dix, D. J.; Houck, K. A.; Martin, M. T.; Richard, A. M.; Setzer, R. W.; Kavlock, R. J. The ToxCast Program for Prioritizing Toxicity Testing of Environmental Chemicals. *Toxicological Sciences* **2007**, *95* (1), 5–12. https://doi.org/10.1093/toxsci/kfl103.
(22) Fröhlich, H.; Speer, N.; Poustka, A.; Beißbarth, T. GOSim – an R-Package for Computation of Information Theoretic GO Similarities between Terms and Gene Products. *BMC Bioinformatics* **2007**, *8* (1), 166. https://doi.org/10.1186/1471-2105-8-166.
(23) Knapen, D.; Angrish, M. M.; Fortin, M. C.; Katsiadaki, I.; Leonard, M.; Margiotta-Casaluci, L.; Munn, S.; O'Brien, J. M.; Pollesch, N.; Smith, L. C.; Zhang, X.; Villeneuve, D. L. Adverse Outcome Pathway Networks I: Development and Applications. *Environmental Toxicology and Chemistry* **2018**, *37* (6), 1723–1733. https://doi.org/10.1002/etc.4125.
(24) Aguayo-Orozco, A.; Audouze, K.; Siggaard, T.; Barouki, R.; Brunak, S.; Taboureau, O. sAOP: Linking Chemical Stressors to Adverse Outcomes Pathway Networks. *Bioinformatics* **2019**, *35* (24), 5391–5392. https://doi.org/10.1093/bioinformatics/btz570.
(25) Jeong, J.; Choi, J. Adverse Outcome Pathways Potentially Related to Hazard Identification of Microplastics Based on Toxicity Mechanisms. *Chemosphere* **2019**, *231*, 249–255. https://doi.org/10.1016/j.chemosphere.2019.05.003.

### Supporting Figures

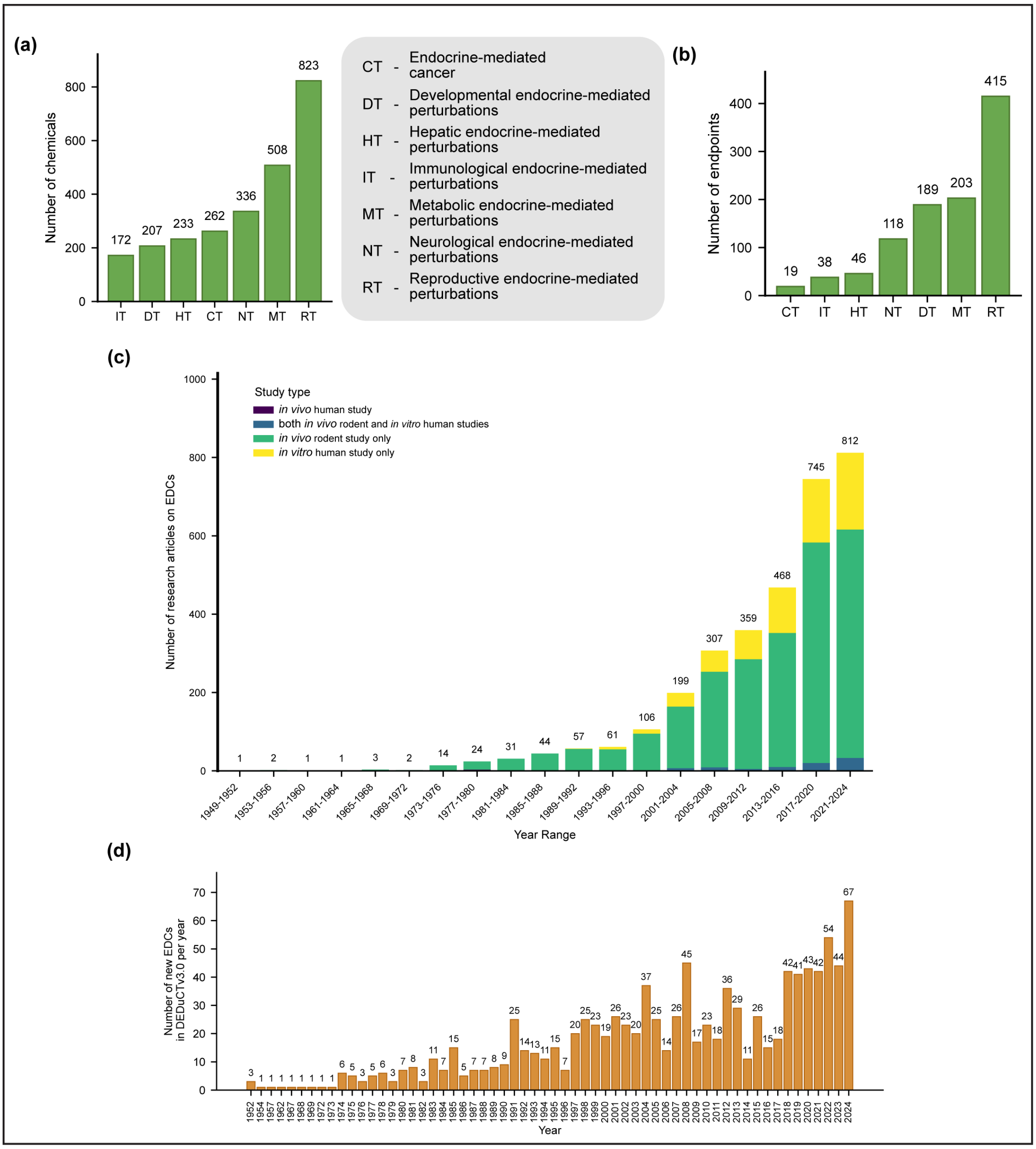


**Figure S1.** (**a**) The distribution of the 796 endocrine-mediated endpoints in DEDuCTv3.0 into the seven systems-level perturbation categories. (**b**) The distribution of chemicals within DEDuCTv3.0 into the seven systems-level perturbation events based on their associated endocrine-mediated endpoint obtained from published literature. (**c**) A chronological analysis of the 3269 published articles supporting 1043 EDCs in DEDuCTv3.0. Note, 32 articles published in January of 2025 are not included. (**d**) Bar plot depicting number of new EDCs identified per year based on published literature compiled in DEDuCTv3.0. Note, 4 new EDCs from January 2025 are not included.

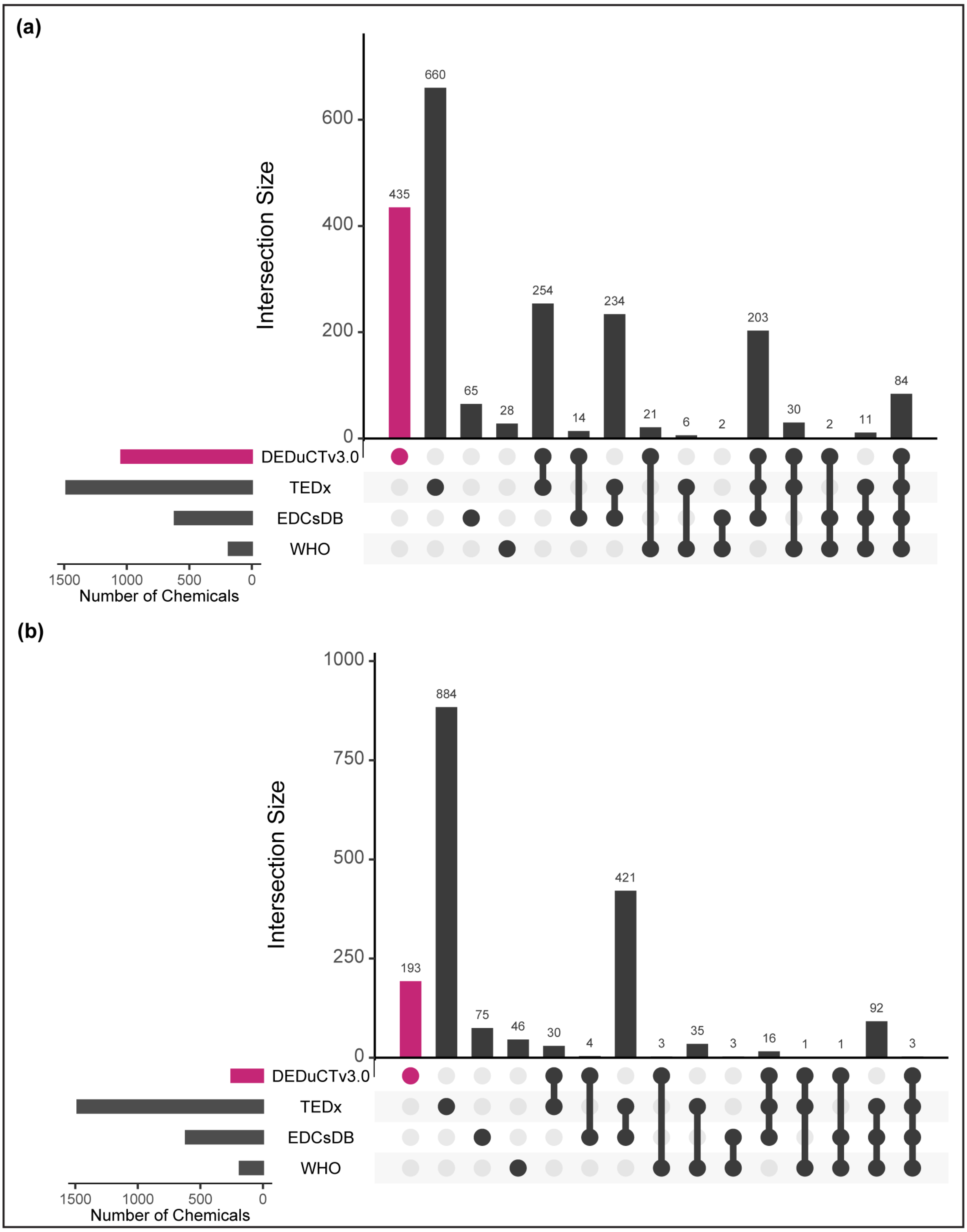


**Figure S2.** UpSet plot representation of unique overlaps between EDCs in DEDuCTv3.0 and other EDC-relevant databases such as the Endocrine Disruption Exchange (TEDx), EDCs Databank version 2015 (EDCsDB) and World Health Organization (WHO) report. (**a**) Comparison with all 1043 EDCs compiled in DEDuCTv3.0. (**b**) Comparison with the 251 new EDCs compiled in DEDuCTv3.0. These UpSet plots were generated using UpSetR package in R.

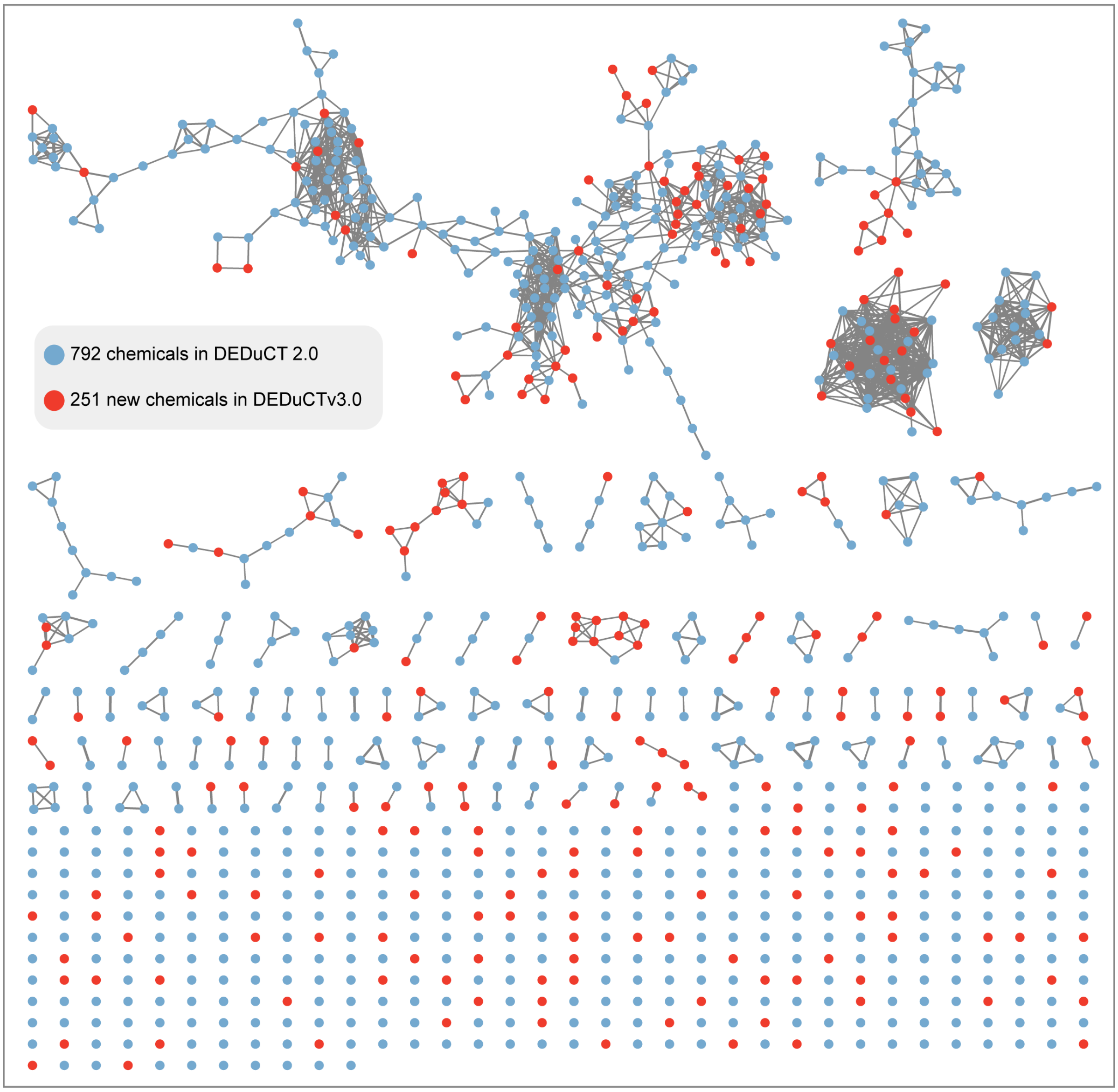


**Figure S3.** Network visualization of the chemical similarity network (CSN) constructed based on the Tanimoto coefficients computed for each pair of chemicals using ECFP4 fingerprints. In this figure, edges are filtered to represent only those with Tanimoto coefficient > 0.5, with the edge thickness proportionally scaled to the corresponding values. The chemicals compiled in DEDuCT 2.0 are represented in blue, while the 251 new chemicals curated in DEDuCTv3.0 are shown in red.

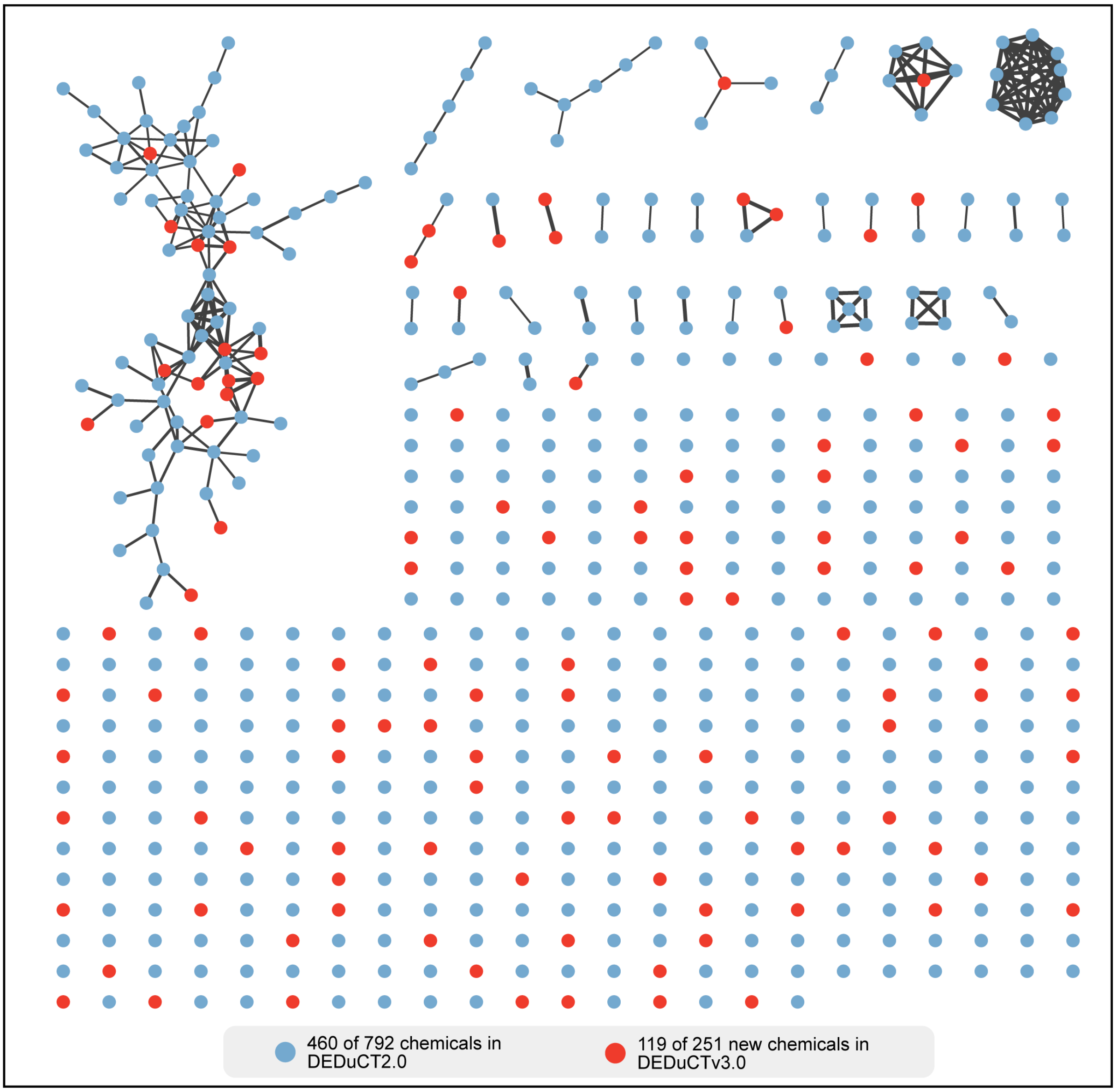


**Figure S4.** Network visualization of the target similarity network (CSN) constructed based on the Jaccard indices computed for each pair of chemicals using information on gene targets. In this figure, edges are filtered to represent only those with Jaccard index > 0.5, with the edge thickness proportionally scaled to the corresponding values. The 460 of 792 chemicals compiled in DEDuCT 2.0 are represented in blue, while the 119 of 251 new chemicals curated in DEDuCTv3.0 are shown in red.

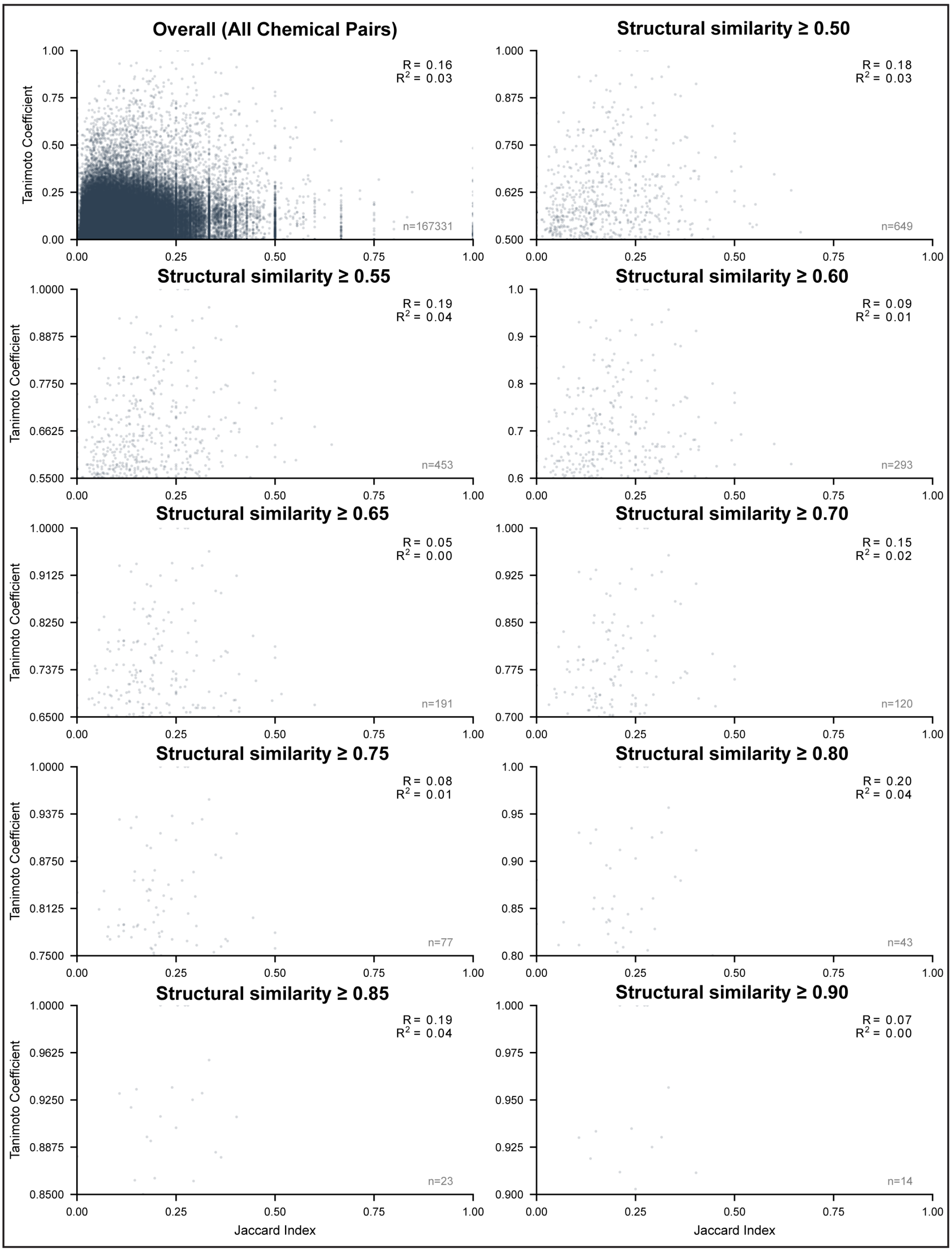


**Figure S5**. The scatter plot of target similarity vs structure similarity between pairs of EDCs. Here, the structure similarity is given by Tanimoto coefficient, and target similarity is given by Jaccard Index. The plots are further divided based on different structural similarity cut-off values. The Pearson correlation coefficient (R) and the coefficient of determination ($R^2$) values are denoted at the top-right of each graph. Further, the number of chemical pairs in each graph is denoted at the bottom right.

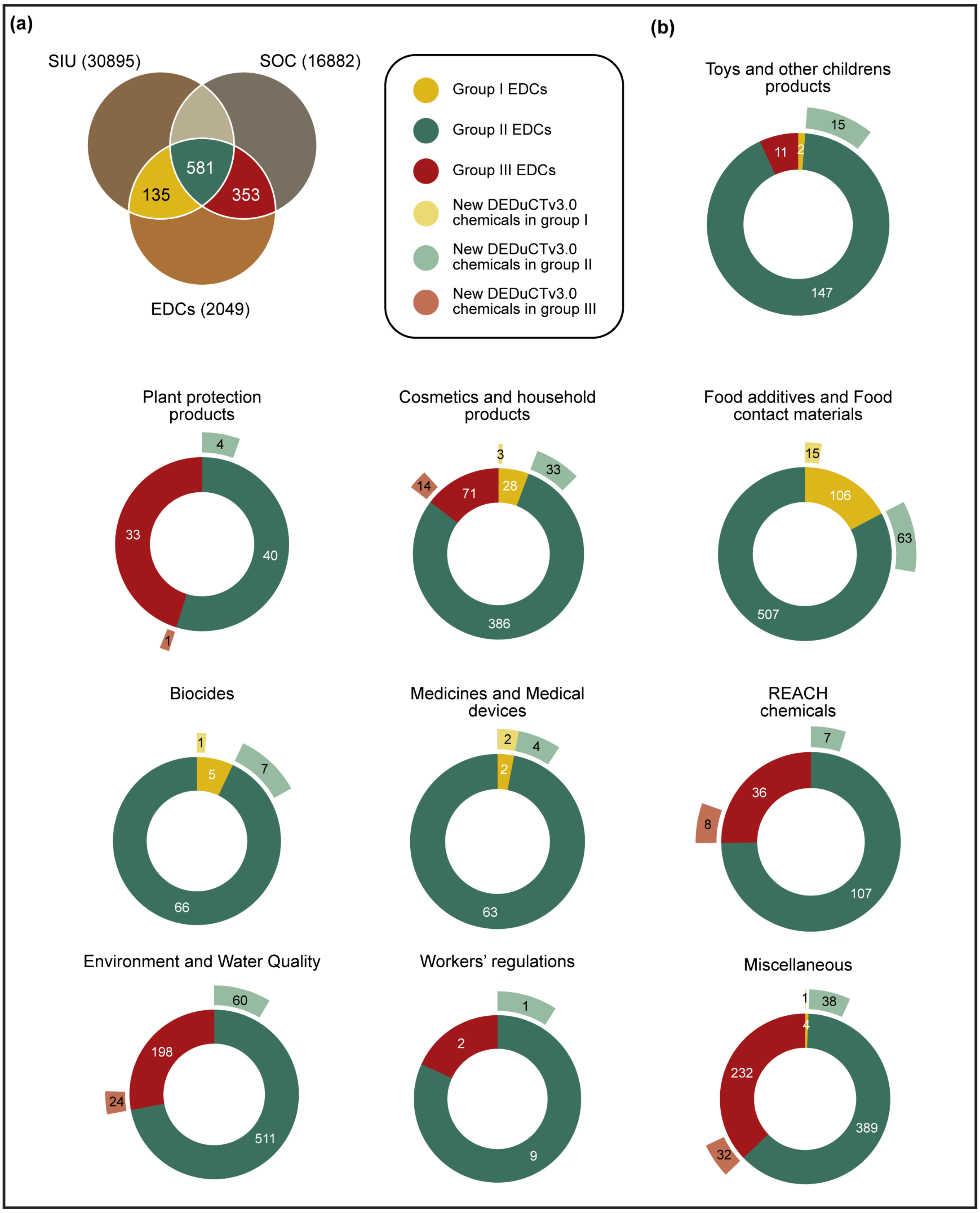


**Figure S6.** Distribution of 2049 unique potential EDCs from DEDuCTv3.0, TEDx, EDCs Databank, and WHO report across 54 chemical lists. **(a)** Venn diagram depicting chemicals categorized into groups I, II, and III. **(b)** Sunburst plots depicting distribution of potential EDCs across 10 categories of chemical lists. Within each category, the inner ring gives the number of potential EDCs categorized as groups I, II, or III, and the outer ring depicts the number of newly compiled chemicals within DEDuCTv3.0 present in each group.

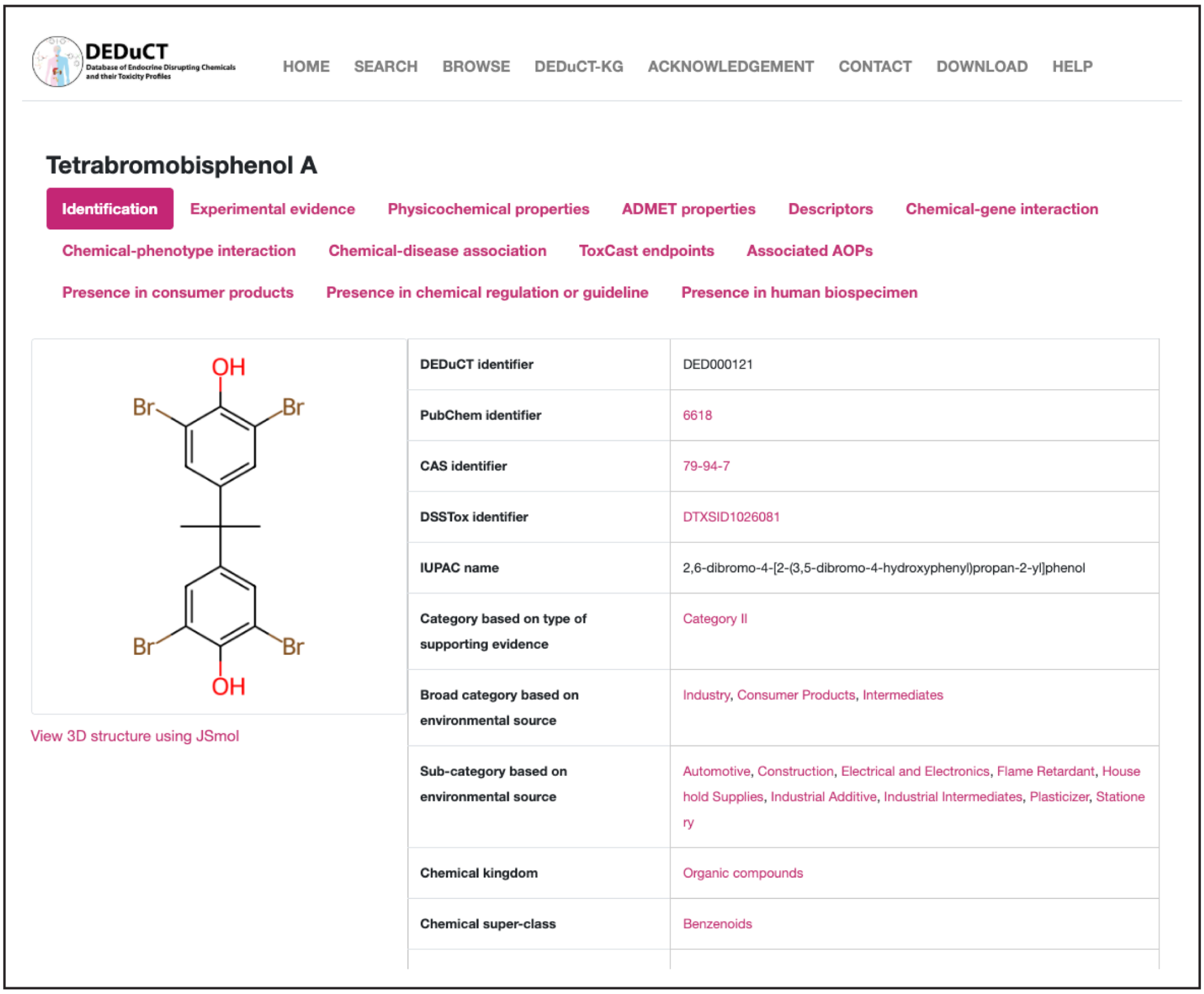


**Figure S7.** Screenshot for the updated chemical information page in DEDuCTv3.0 webserver.

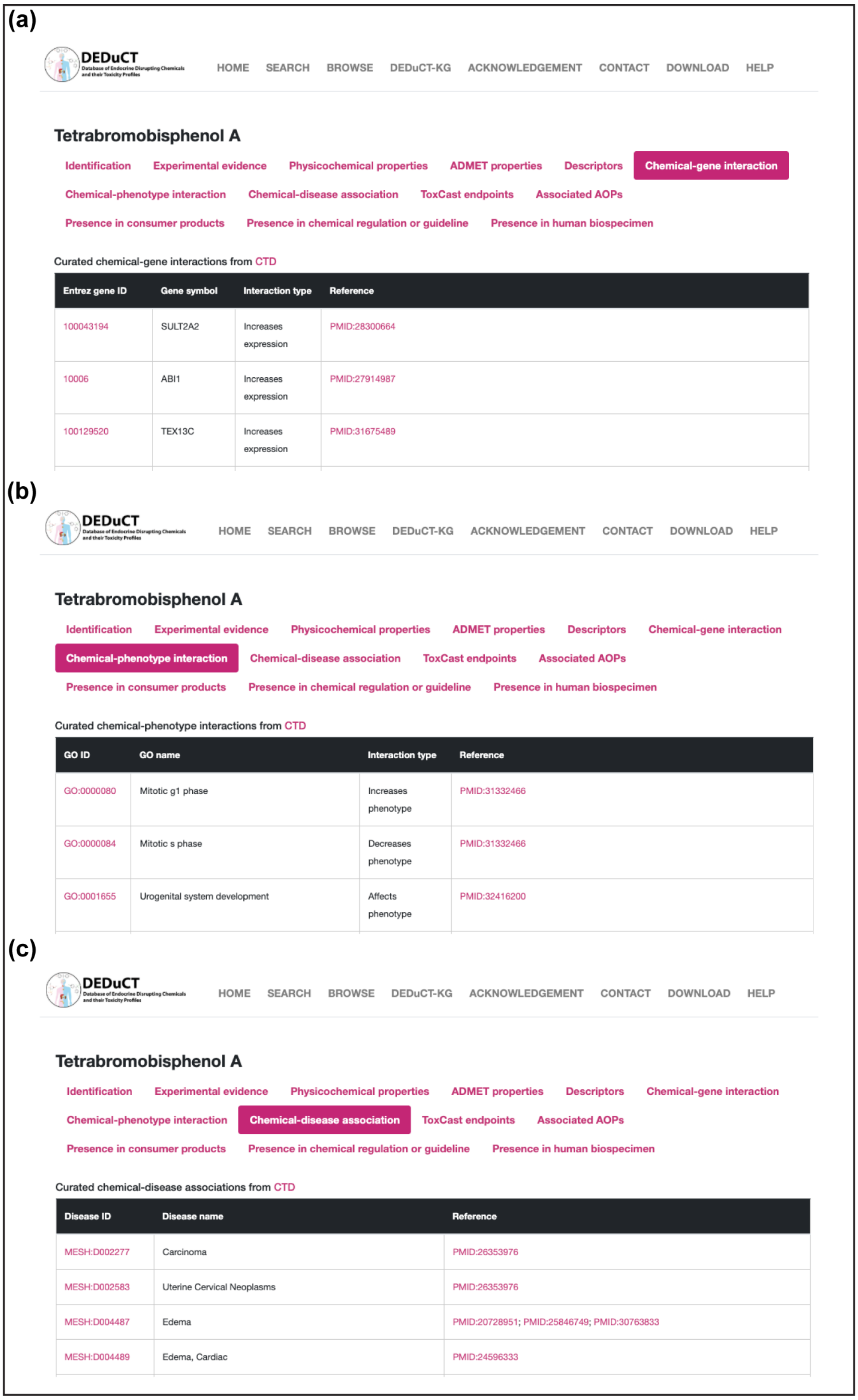


**Figure S8.** Screenshots from DEDuCTv3.0 webserver of the chemical information pages on: (**a**) chemical-gene interactions; (**b**) chemical-phenotype interactions; (**c**) chemical-disease associations.

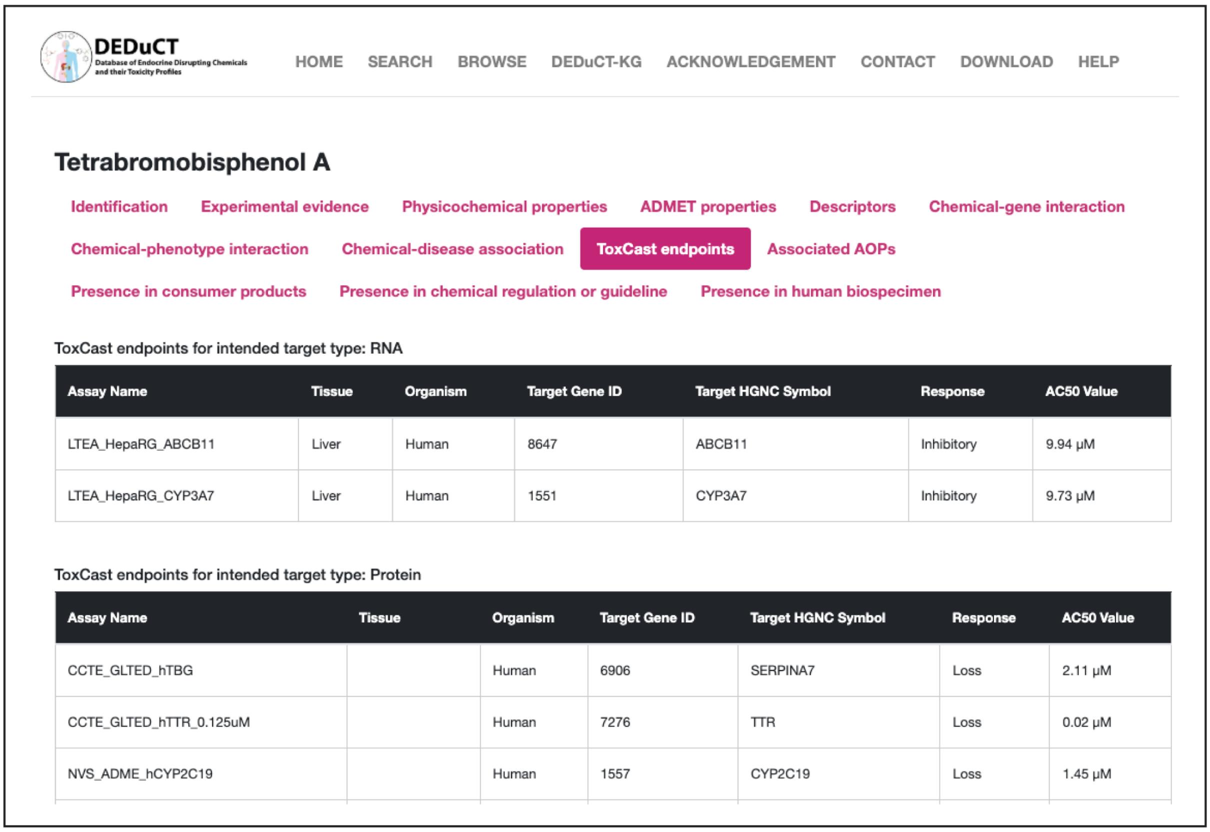


**Figure S9.** Screenshot from DEDuCTv3.0 webserver of the chemical information page on associated ToxCast endpoints.

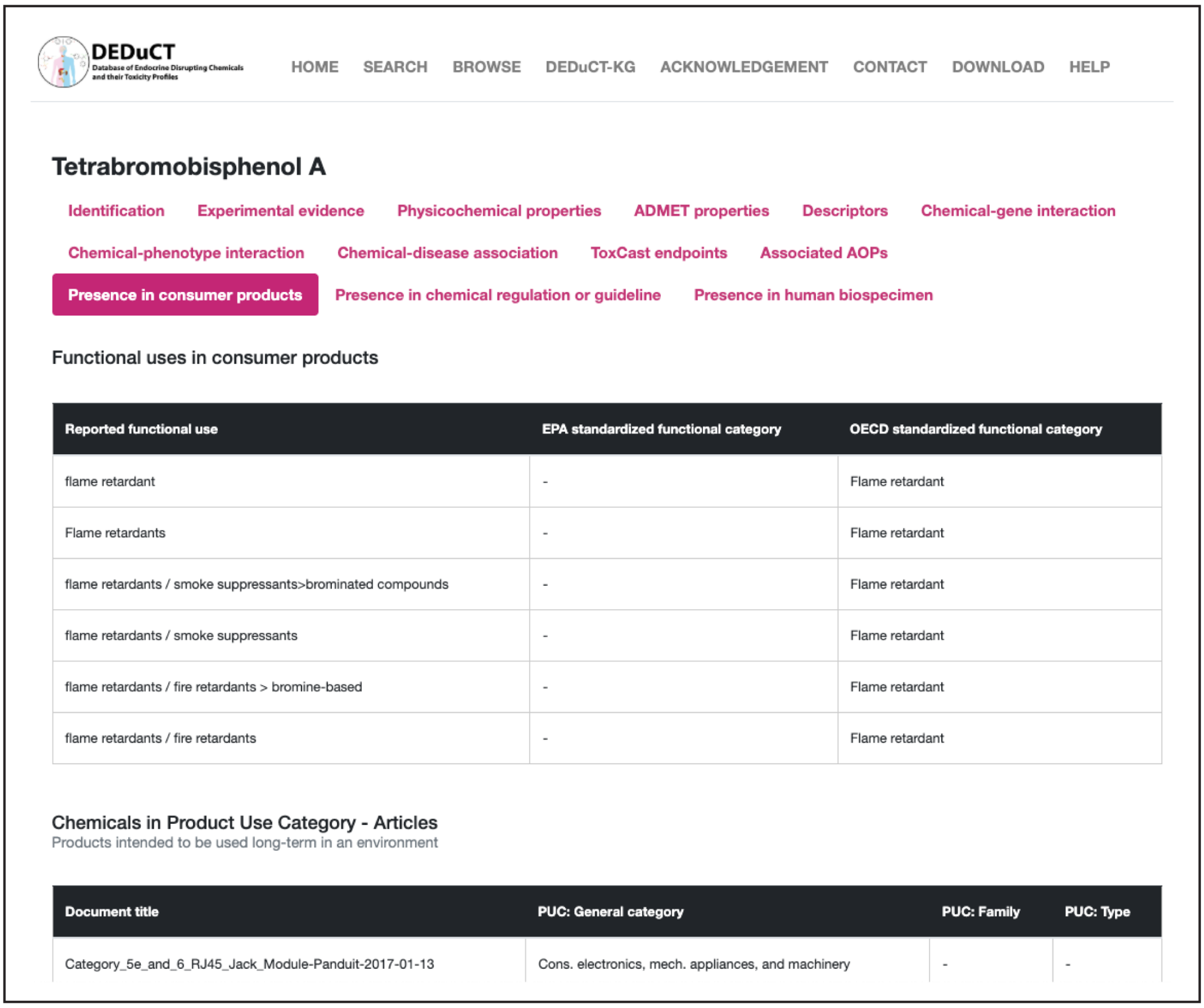


**Figure S10.** Screenshot from DEDuCTv3.0 webserver of the chemical information page on associated data from Chemical and Products Database (CPDat).

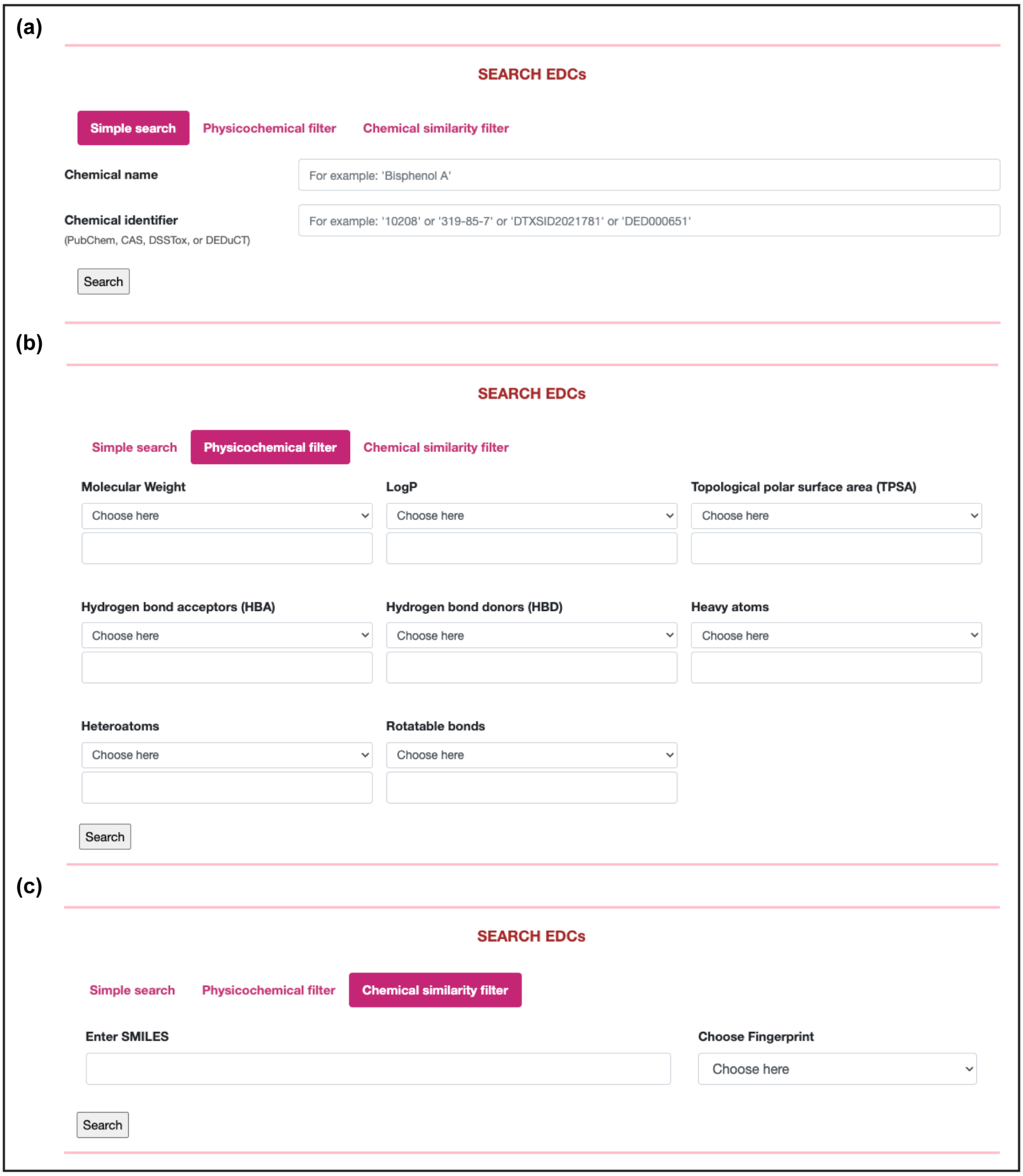


**Figure S11.** Screenshots from DEDuCTv3.0 webserver for the different search options. (**a**) Search based on chemical information such as names, synonyms, PubChem identifier, CASRN, DSSTox identifier, and DEDuCT identifier. (**b**) Search based on physicochemical properties of chemicals. (**c**) Search based on structural similarity.
